# A crosstalk between hepcidin and IRE/IRP pathways controls ferroportin expression and determines serum iron levels in mice

**DOI:** 10.1101/2021.10.29.466457

**Authors:** Edouard Charlebois, Carine Fillebeen, Angeliki Katsarou, Aleksandr Rabinovich, Kazimierz Wisniewski, Vivek Venkataramani, Bernhard Michalke, Anastasia Velentza, Kostas Pantopoulos

**Affiliations:** Lady Davis Institute for Medical Research, Jewish General Hospital and Department of Medicine, McGill University, Montreal, Quebec, Canada; Ferring Research Institute Inc, San Diego, CA; Department of Medicine II, Hematology/Oncology, University Hospital Frankfurt, Frankfurt, Germany; Institute of Pathology, University Medical Center Göttingen (UMG), Göttingen, Germany; Helmholtz Zentrum München GmbH – German Research Center for Environmental Health, Research Unit Analytical BioGeoChemistry, Neuherberg, Germany

## Abstract

The iron hormone hepcidin is transcriptionally activated by iron or inflammation via distinct, partially overlapping pathways. We addressed how iron affects inflammatory hepcidin levels and the ensuing hypoferremic response. Dietary iron overload did not mitigate hepcidin induction in LPS-treated wt mice but prevented effective inflammatory hypoferremia. Likewise, LPS modestly decreased serum iron in hepcidin-deficient Hjv-/-mice, model of hemochromatosis. Synthetic hepcidin triggered hypoferremia in control but not iron-loaded wt animals. Furthermore, it dramatically decreased hepatic and splenic ferroportin in Hjv-/-mice on standard or iron-deficient diet, but only triggered hypoferremia in the latter. Mechanistically, iron antagonized hepcidin responsiveness by inactivating IRPs in the liver and spleen, to stimulate ferroportin mRNA translation. Prolonged LPS treatment eliminating ferroportin mRNA permitted hepcidin-mediated hypoferremia in iron-loaded mice. Thus, *de novo* ferroportin synthesis is critical determinant of serum iron and finetunes hepcidin-dependent functional outcomes. Our data uncover a crosstalk between hepcidin and IRE/IRP systems that controls tissue ferroportin expression and determines serum iron levels. Moreover, they suggest that hepcidin supplementation therapy is more efficient combined with iron depletion.

## Introduction

Systemic iron balance is controlled by hepcidin a peptide hormone that is produced by hepatocytes in the liver and operates in target cells by binding to the iron exporter ferroportin (1, 2). This results in ferroportin internalization and lysosomal degradation but also directly inhibits ferroportin function by occluding its iron export channel (3, 4). Ferroportin is highly expressed in duodenal enterocytes and tissue macrophages, which are instrumental for dietary iron absorption and iron recycling from senescent erythrocytes, respectively. Ferroportin is also expressed in hepatocytes, where excess iron is stored and can be mobilized on demand. Hepcidin-mediated ferroportin inactivation inhibits iron entry into plasma. This is a critical homeostatic response against iron overload, but also an innate immune response against infection (5). Thus, hepcidin expression is induced when systemic iron levels are high to prevent dietary iron absorption, or under inflammatory conditions to promote iron retention within ferroportin-expressing cells and render the metal unavailable to extracellular pathogens.

The hepcidin-encoding *HAMP* gene is transcriptionally induced by iron or inflammatory stimuli via BMP/SMAD (6) or IL-6/STAT3 (7) signaling, respectively. These pathways crosstalk at different levels. For instance, the BMP co-receptor hemojuvelin (HJV), a potent enhancer of iron-dependent BMP/SMAD signaling, is also essential for inflammatory induction of hepcidin. Thus, Hjv-/-mice, a model of juvenile hemochromatosis characterized by severe iron overload and hepcidin deficiency (8), exhibit blunted inflammatory induction of hepcidin and fail to mount a hypoferremic response following LPS treatment or infection with *E. coli* (9). Excess iron inhibits hepcidin induction via the BMP/SMAD and IL-6/STAT3 signaling pathways in cultured cells (10, 11), but the *in vivo* relevance of these findings is not known.

Hepcidin-dependent inhibition of ferroportin activity and expression is a major but not the sole contributor to inflammatory hypoferremia (12, 13). This is related to the fact that ferroportin expression is regulated by additional transcriptional and post-transcriptional mechanisms (14). Thus, ferroportin transcription is induced by iron (15) and suppressed by inflammatory signals (16), while translation of *Fpn(+IRE)* mRNA, the major ferroportin transcript that harbors an “iron responsive element” (IRE) within its 5’ untranslated regions (5’ UTR) is controlled by “iron regulatory proteins” (IRPs), IRP1 and IRP2. The IRE/IRP system accounts for coordinate post-transcriptional regulation of iron metabolism proteins in cells (17, 18). In a homeostatic response to iron deficiency, IRPs bind to the IRE within the *Fpn(+IRE)* and ferritin (*Fth1* and *Ftl1*) mRNAs, inhibiting their translation. IRE/IRP interactions do not take place in iron-loaded cells, allowing *de novo* ferroportin and ferritin synthesis to promote iron efflux and storage, respectively. The impact of the IRE/IRP system on regulation of tissue ferroportin and serum iron is not well understood.

The aim of this work was to elucidate mechanisms by which systemic iron overload affects hepcidin expression and downstream responses, especially under inflammatory conditions. Utilizing wild type and Hjv-/-mice, we demonstrate that serum iron levels reflect regulation of ferroportin in the liver and spleen by multiple signals. We further show that effective hepcidin-mediated hypoferremia is antagonized by compensatory mechanisms aiming to prevent cellular iron overload. Our data uncover a crosstalk between hepcidin and the IRE/IRP system that controls ferroportin expression in the liver and spleen, and thereby determines serum iron levels.

## Results

### Dietary iron overload does not prevent further inflammatory Hamp mRNA induction in LPS-treated wt mice, but mitigates hepcidin responsiveness

In an exploratory experiment, wt mice were subjected to dietary iron loading by feeding a high-iron diet (HID) for short (1 day), intermediate (1 week) or long (5 weeks) time intervals; control animals remained on standard diet (SD). As expected, mice on HID for 1 day manifested maximal increases in serum iron (Fig. S1A) and transferrin saturation (Fig. S1B). They retained physiological liver iron content (LIC; Fig. S1C) and serum ferritin (Fig. S1D), a reflection of LIC. Serum iron and transferrin saturation plateaued after longer HID intake, while LIC and serum ferritin gradually increased to peak at 5 weeks. The dietary iron loading promoted gradual upregulation of serum hepcidin (Fig. S1E) and liver *Hamp* mRNA (Fig. S1F), with highest values at 5 weeks. This could not prevent chronic dietary iron overload, in agreement with earlier findings (19, 20).

LPS triggered appropriate hepcidin induction and a robust hypoferremic response in control mice. Interestingly, LPS-induced inflammation resulted in further proportional increase in hepcidin and *Hamp* mRNA in dietary iron-loaded mice (Fig. S1E-F). This was accompanied by significant drops in serum iron and transferrin saturation (Fig. S1A-B). However, values did not reach the nadir of LPS-treated control animals and were increasing in mice on HID for longer periods, despite significant hepcidin accumulation. These data suggest that hepatic iron overload does not prevent inflammatory induction of hepcidin; however, it impairs its capacity to decrease serum iron.

### Uncoupling inflammatory hepcidin induction from hypoferremic response in wt and Hjv-/- mice following dietary iron manipulations

To further explore the potential of hepcidin to promote hypoferremia under iron overload, wt and Hjv-/-mice, a model of hemochromatosis, were subjected to dietary iron manipulations. Wt mice were fed SD or HID, and Hjv-/-mice were fed SD or an iron-deficient diet (IDD) for 5 weeks, to achieve a broad spectrum of hepcidin regulation. Wt mice on HID and Hjv-/-mice on SD or IDD manifested similarly high serum iron and transferrin saturation (Fig. 1A-B). Serum non-transferrin bound iron (NTBI) levels appeared modestly elevated in the dietary and genetic iron overload models and seemed to decrease in Hjv-/-mice following IDD intake (Fig. 1C). LIC was substantially reduced in Hjv-/-mice in response to IDD, but also compared to wt mice on HID (Fig. 1D). The quantitative LIC data were corroborated histologically by Perls staining (Fig. 1E and S2A). Dietary iron loading increased splenic iron in wt mice and confirmed that Hjv-/-mice fail to retain iron in splenic macrophages (Fig. S2B). As expected, serum hepcidin (Fig. 1F) and liver *Hamp* mRNA (Fig. 1G) were maximally induced in HID-fed wt mice and were low in Hjv-/-mice on SD, and further suppressed to undetectable levels following IDD intake.

**Fig. 1.**
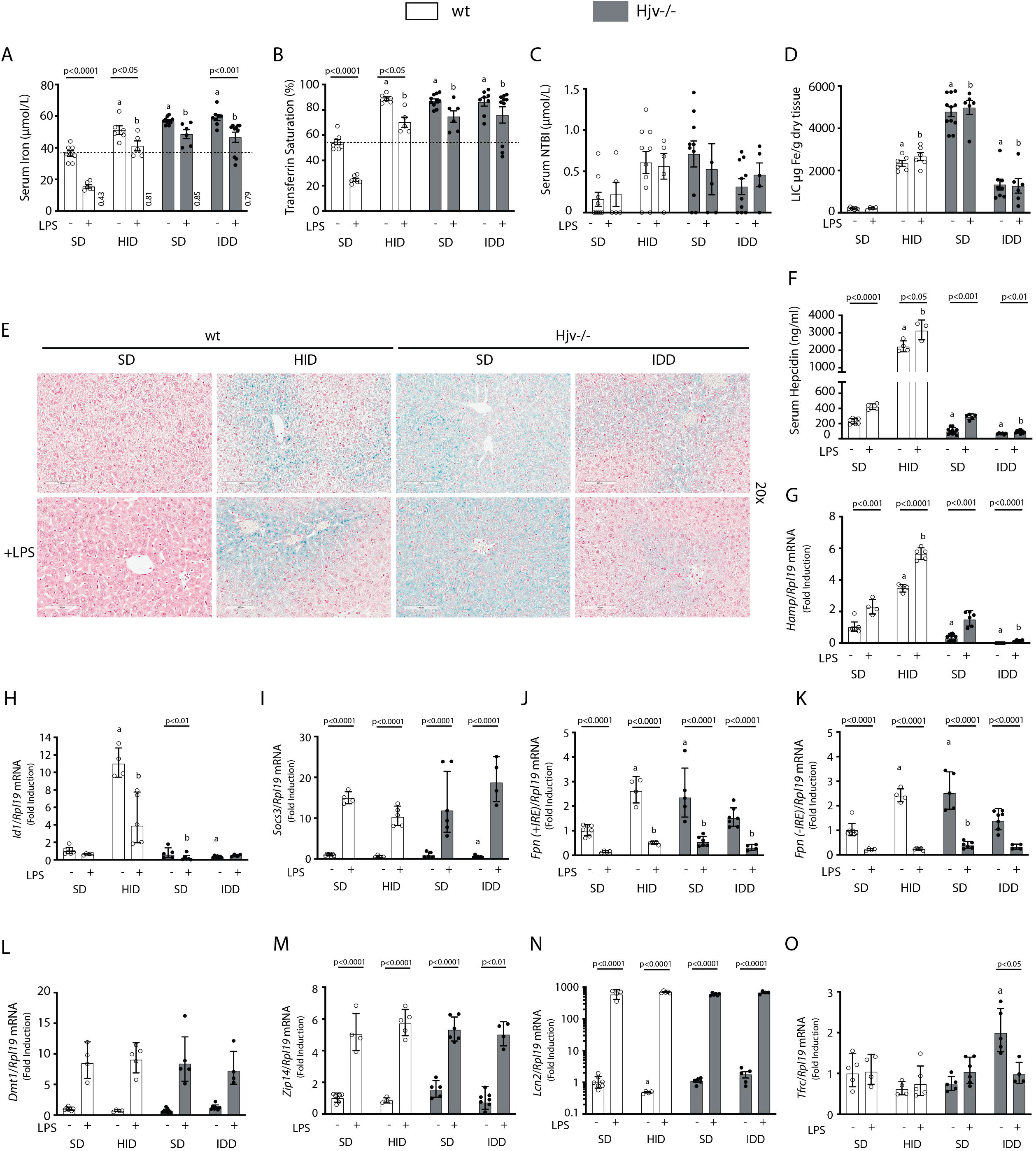
Iron overload blunts hepcidin responsiveness to LPS-induced inflammation. Four-week-old male wt mice (n=12-14 per group) were placed on HID for five weeks. Conversely, age- and sex-matched isogenic Hjv-/-mice (n=12-14 per group) were placed on IDD for five weeks to prevent excessive iron overload. Controls from both genotypes were kept on SD. Half of the mice were injected with saline and the other half with 1 µg/g LPS; all animals were sacrificed 4 hours later. Sera were collected by cardiac puncture and analyzed for: (A) iron, (B) transferrin saturation, (C) NTBI, and (F) hepcidin. Livers were dissected and processed for LIC quantification by the ferrozine assay (D) and for histological detection of iron deposits by Perls’ staining (E; magnification: 20x). Livers were also used for qPCR analysis of following mRNAs:(G) *Hamp*, (H) Id1, (I) Socs3, (J) *Fpn(+IRE)*, (K) *Fpn(-IRE)*, (L) *Dmt1*, (M) *Zip14*, (N) *Lcn2* and (O) *Tfrc*. The dotted line in (A) and (B) indicates baseline serum iron and transferrin saturation, respectively, of control wt mice on SD. Values in (A) represent ratios of serum iron levels between untreated and LPS-treated mice. Data in (A-F) are presented as the mean±SEM while in (G-O) are presented as geometric mean±SD. Statistically significant differences (p<0.05) compared to values from saline- or LPS-treated wt control mice are indicated by a or b, respectively.

LPS reduced serum iron and transferrin saturation in hyperferremic wt mice on HID and Hjv-/-mice on SD or IDD, but not below baseline of control wt mice on SD, the only animals that developed a robust hypoferremic response (Fig. 1A-B); see also ratios of serum iron levels between untreated and LPS-treated mice in Fig. 1A. The LPS treatment was associated with significant accumulation of hepcidin (Fig. 1F) and induction of *Hamp* mRNA (Fig. 1G) in all experimental groups, while NTBI (Fig. 1C) and LIC (Fig. 1D) were unaffected. Notably, LPS-treated wt mice on HID and Hjv-/-mice on IDD exhibited dramatic differences in *Hamp* mRNA but similar blunted hypoferremic response to the acute inflammatory stimulus. Thus, the profound hepcidin induction in iron-loaded wt mice cannot decrease serum iron below that of iron-depleted Hjv-/-mice with negligible hepcidin, which indicates reduced hepcidin responsiveness. In support to this interpretation, *Id1* and *Socs3* mRNAs (Fig. 1H-I), which are markers of BMP/SMAD and IL-6/STAT3 signaling, respectively, were appropriately induced by dietary iron loading or LPS treatment in wt mice. Thus, the major hepcidin signaling pathways were intact under these experimental conditions.

Serum iron levels are also controlled by hepcidin-independent mechanisms (12, 13). To explore their possible contribution in our experimental setting, we analyzed expression of genes involved in iron transport in the liver, an organ that contributes to iron sequestration during inflammation. Ferroportin is encoded by two alternatively spliced transcripts, *Fpn(+IRE)* and *Fpn(-IRE)* (21). Both of them were significantly increased in the liver of iron-loaded wt mice on HID and Hjv-/-mice on SD, which is consistent with transcriptional induction (15), and were strongly suppressed by LPS (Fig. 1J-K). The LPS treatment induced *Dmt1, Zip14* and *Lcn2* mRNAs in all animals (Fig. 1L-N). These encode the divalent metal transporter DMT1, the NTBI transporter Zip14 and the siderophore-binding protein Lcn2, respectively; *Lcn2* mRNA induction was dramatic. The transferrin receptor 1-encoding *Tfrc* mRNA was largely unaffected by LPS, except for a reduction in Hjv-/-mice on IDD (Fig. 1O). The above data indicate that LPS-induced inflammation triggers transcriptional responses favoring reduced iron efflux from the liver and increased uptake of NTBI by liver cells.

To assess the downstream function of hepcidin, we analyzed tissue ferroportin levels. Immunohistochemical staining of liver sections revealed strong ferroportin expression in Kupffer cells, predominantly in periportal areas, under all experimental conditions (Figs. 2A and S3). Hepatocellular ferroportin staining is also evident in the iron overload models, mostly in periportal hepatocytes (Fig. S3), and in line with recent data (22). LPS triggered redistribution and decreased expression of ferroportin in Kupffer cells from wt but not Hjv-/-mice (Fig. S3), as reported (9).

**Fig. 2.**
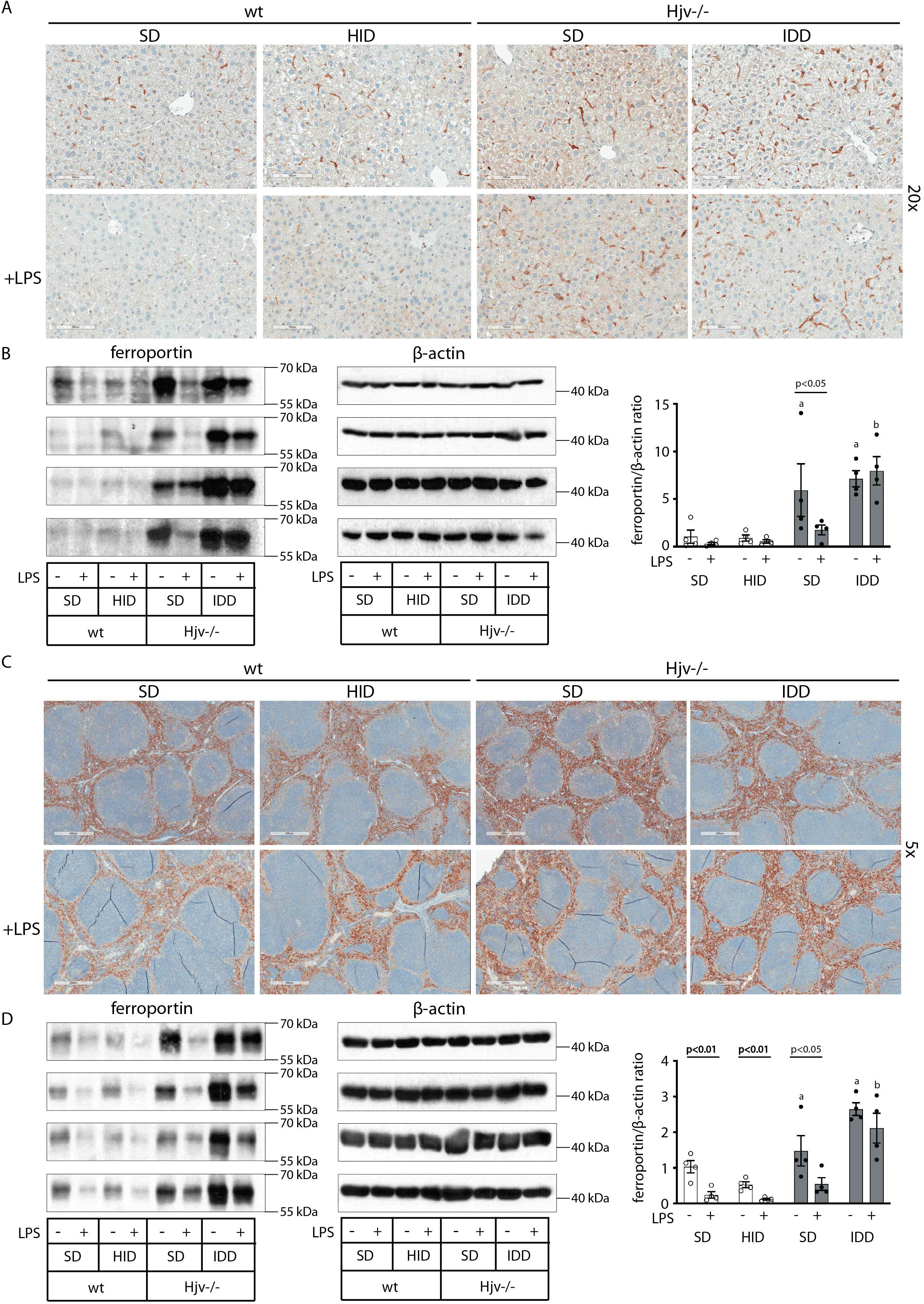
Effects of LPS on hepatic and splenic ferroportin of iron-manipulated wt and Hjv-/-mice. Livers and spleens from mice described in Fig. 1 were dissected and processed for immunohistochemical and biochemical analysis of ferroportin. Immunohistochemical staining of ferroportin in liver (A) and spleen (C) sections (magnification for liver is 20x and for spleen 5x). Western blot for ferroportin and β-actin in liver (B) and spleen (D) extracts from four representative mice in each condition. Blots were quantified by densitometry and ferroportin/β-actin ratios are shown on the right. Densitometric data are presented as the mean±SEM. Statistically significant differences (p<0.05) compared to values from saline- or LPS-treated wt control mice are indicated by a or b, respectively. Statistics in bold were performed using unpaired Student’s t test.

We further analyzed ferroportin in liver homogenates by Western blotting. Levels of biochemically detectable liver ferroportin differed substantially between wt and Hjv-/-mice. Thus, they were relatively low in the former and highly induced in the latter (Fig. 2B), independently of iron load. The differences were more dramatic compared to those observed by immunohistochemistry (Figs. 2A and S3). Conceivably, the strong ferroportin signal in Western blots from Hjv-/-liver homogenates reflects high ferroportin expression in hepatocytes, which are the predominant cell population and make up ∼80% of the liver cell mass (23). Yet, hepatocellular ferroportin is less visible by immunohistochemistry because the signal is substantially weaker compared to that in Kupffer cells (see also Fig. 5E). Interestingly, the LPS treatment visibly suppressed total liver ferroportin in Hjv-/-mice on SD but not IDD, and appeared to modestly reduce it in wt mice (Fig. 2B); albeit, without statistical significance. These data are consistent with negative regulation of ferroportin by residual LPS-induced hepcidin in Hjv-/-mice on SD, which could explain the small drop in serum iron and transferrin saturation under these acute inflammatory conditions, as reported (9). However, liver ferroportin remained detectable and apparently functional, as it did not allow significant iron sequestration and dramatic drop in serum iron. Notably, persistence of relatively high serum iron is also evident in LPS-treated wt mice on HID, despite maximal hepcidin and minimal liver ferroportin levels.

Next, we analyzed ferroportin in the spleen, an organ with erythrophagocytic macrophages that plays an important role in body iron traffic (24). Immunohistochemical analysis shows that LPS reduced ferroportin in red pulp splenic macrophages from wt mice on SD, but this effect was less evident in wt mice on HID and in Hjv-/-mice on SD or IDD (Figs. 2C and S4). Western blot analysis shows a stronger ferroportin signal in splenic extracts from Hjv-/-animals (Fig. 2D), consistent with immunohistochemistry. However, in this assay LPS suppressed splenic ferroportin in wt animals and in Hjv-/-mice on SD, but not IDD. This could be a result of residual hepcidin upregulation (Fig. 1F-G), while the lack of significant splenic ferroportin suppression in Hjv-/-mice on IDD may denote hepcidin insufficiency. In any case, the relatively high circulating iron levels in dietary iron-loaded and LPS-treated wt mice indicates continuous iron efflux to plasma despite hepcidin excess.

### Insufficient hepcidin leads to blunted hypoferremic response in iron overload

We used human synthetic hepcidin to address whether the failure of mouse models of iron overload to mount an appropriate hypoferremic response to acute inflammation is caused by endogenous hepcidin insufficiency or other mechanisms. Wt and Hjv-/-mice subjected to dietary iron manipulations received 2.5 μg/g synthetic hepcidin every two hours for a total of four intraperitoneal injections. Each dose corresponds to ∼200-fold excess over endogenous circulating hepcidin in control wt animals. The treatment caused hypoferremia in wt mice on SD but not HID, where the decrease in serum iron was significant but well above baseline of untreated wt controls (Fig. 3A-B); see also ratios of serum iron levels between untreated and hepcidin-treated mice in Fig. 3A. Likewise, synthetic hepcidin significantly decreased serum iron but failed to cause dramatic hypoferremia in hepcidin-deficient Hjv-/-mice on SD. Notably, hepcidin administration was much more effective in relatively iron-depleted Hjv-/-mice on IDD, and lowered serum iron and transferrin saturation below baseline. The treatments significantly reduced NTBI in Hjv-/-mice on SD, with a trend in mice on IDD (Fig. 3C) but did not affect LIC or splenic iron content (SIC) under any experimental conditions (Figs. 3D-E and S5). Serum iron represents <2% of total tissue iron and therefore its acute fluctuations are not expected to dramatically alter LIC or SIC.

**Fig. 3.**
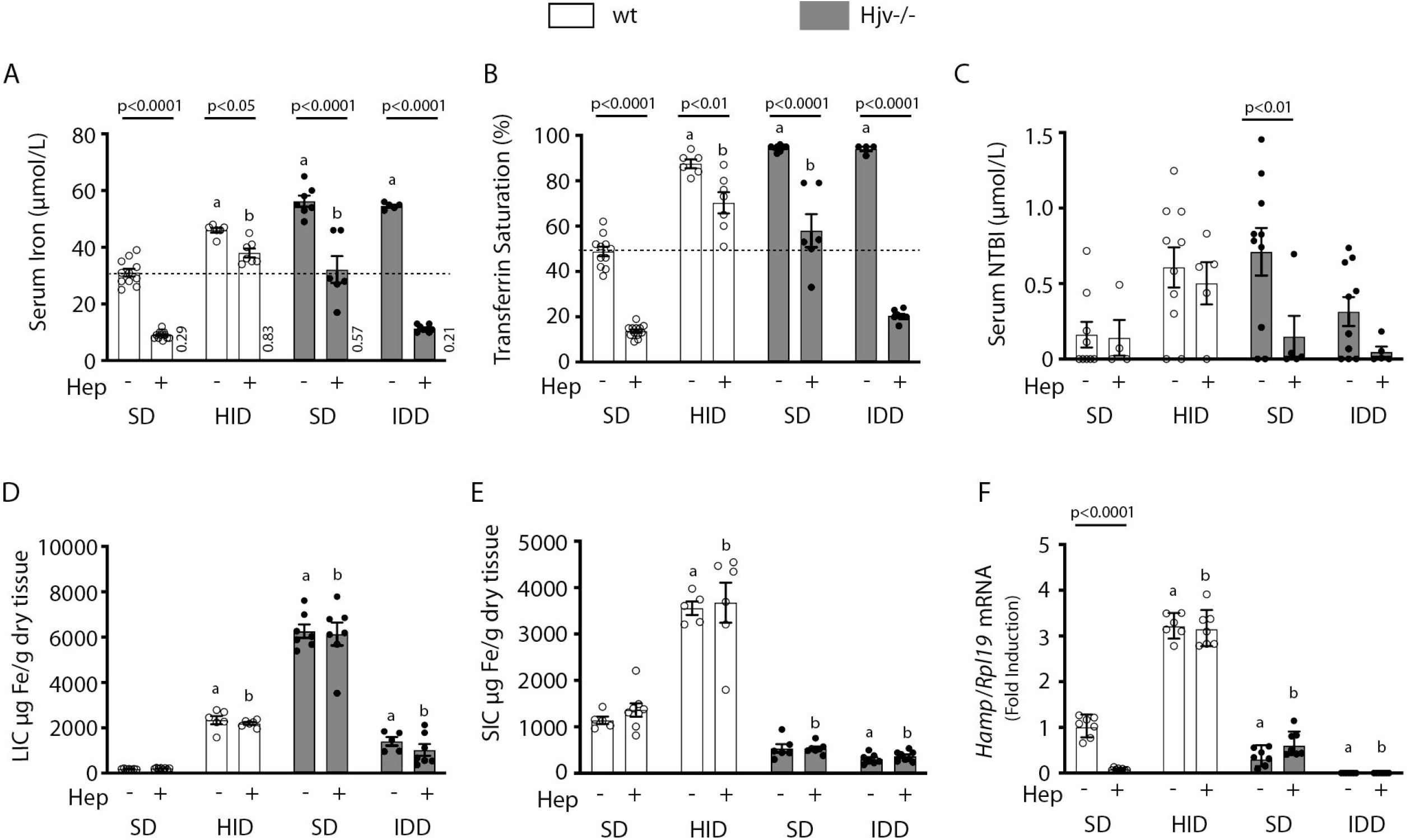
Iron depletion of Hjv-/-mice improves the efficacy of synthetic hepcidin to promote hypoferremia. Four-week-old wt male mice (n=12-14 per group) were placed on HID for five weeks. Conversely, age- and sex-matched isogenic Hjv-/-mice (n=12-14 per group) were placed on IDD for five weeks to prevent excessive iron overload. Controls from both genotypes were kept on SD. Half of the mice were injected every 2 hours for a total of 4 injections with saline, and the other half with 2.5 µg/g synthetic hepcidin. Sera were collected by cardiac puncture and analyzed for: (A) iron, (B) transferrin saturation, and (C) NTBI. Livers and spleens were dissected and processed for analysis of: (D) LIC and (E) SIC by the ferrozine assay. (F) qPCR analysis of liver *Hamp* mRNA. The dotted line in (A) and (B) indicates baseline serum iron and transferrin saturation, respectively, of control wt mice on SD. Values in (A) represent ratios of serum iron levels between untreated and hepcidin-treated mice. Data in (A-E) are presented as the mean±SEM and in (F) as geometric mean±SD. Statistically significant differences (p<0.05) compared to values from saline- or hepcidin-treated wt control mice are indicated by a or b, respectively.

Synthetic hepcidin led to significant reduction of endogenous *Hamp* mRNA in wt mice on SD (Fig. 3F), as earlier reported (25). Conceivably, this is related to destabilization of the Hamp inducer Tfr2 in the liver (Fig. S6), a known response to hypoferremia (26). Synthetic hepcidin did not promote inflammation, iron perturbations or alterations in BMP/SMAD signaling in the liver, as judged by the unaltered expression of hepatic *Fpn(+IRE), Socs3, Id1* and *Bmp6* mRNAs (Fig. S7A-D). Moreover, synthetic hepcidin did not affect *Dmt1, Zip14, Lcn2* or *Tfrc* mRNAs (Fig. S7E-H), which encode iron transporters; *Zip14* and *Lcn2* are also inflammatory markers.

Next, we analyzed liver ferroportin by immunohistochemistry. Figs. 4A and S8 show that exogenous hepcidin decreased ferroportin signal intensity in all animal groups to varying degrees. The hepcidin effect was particularly noticeable in hepatocytes from Hjv-/-mice (see low magnification images in Fig. S8). Kupffer cells seemed to retain some ferroportin in all groups except Hjv-/-mice on IDD. Interestingly, while synthetic hepcidin decreased ferroportin signal intensity in Kupffer cells, it did not alter intracellular ferroportin distribution as would be expected based on the data in LPS-treated wt mice (Fig. 4A).

**Fig. 4.**
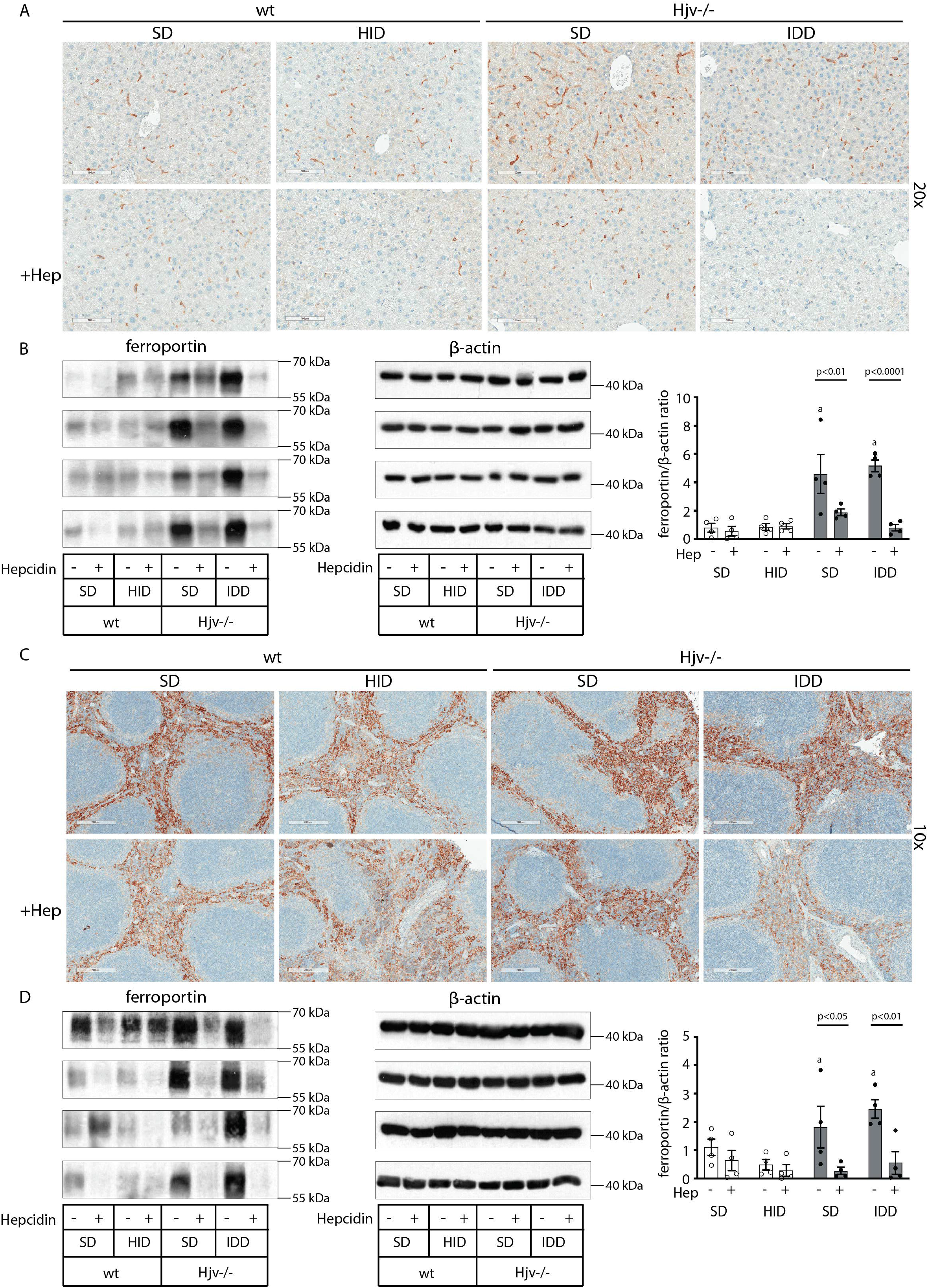
Effects of synthetic hepcidin on hepatic and splenic ferroportin of iron-manipulated wt and Hjv-/-mice. Livers and spleens from mice described in Fig. 3 were dissected and processed for immunohistochemical and biochemical analysis of ferroportin. Immunohistochemical staining of ferroportin in liver (A) and spleen (C) sections (magnification for liver is 20x and for spleen 10x). Western blot for ferroportin and β-actin in liver (B) and spleen (D) extracts from four representative mice in each condition. Blots were quantified by densitometry and ferroportin/β-actin ratios are shown on the right. Densitometric data are presented as the mean±SEM. Statistically significant differences (p<0.05) compared to values from saline- or hepcidin-treated wt control mice are indicated by a or b, respectively.

Western blotting confirmed that total liver ferroportin is highly induced in Hjv-/-mice (Fig. 4B). Again, the signal intensity can be attributed to protein expressed in hepatocytes. The treatment with synthetic hepcidin did not significantly affect liver ferroportin in wt mice (either on SD or HID), but substantially reduced it in Hjv-/-mice, to almost wt levels. The effect appeared more pronounced in Hjv-/-mice on IDD; nevertheless, ferroportin remained detectable.

Splenic ferroportin was reduced in all animal groups following hepcidin treatment, with stronger effects visualized by immunohistochemistry in wt mice on SD and Hjv-/-mice on IDD (Figs. 4C and S9). At the biochemical level, ferroportin expression was again much stronger in the spleen of Hjv-/-mice (Fig. 4D). Synthetic hepcidin did not significantly affect splenic ferroportin in wt mice, but dramatically reduced it in all Hjv-/-mice.

Taken together, our data suggest that synthetic hepcidin overcomes endogenous hepcidin deficiency in Hjv-/-mice. However, it only triggers hypoferremia in these animals following relative iron depletion. On the other hand, in iron-loaded wt mice with already high endogenous hepcidin, excess synthetic hepcidin fails to promote hypoferremia.

### Dietary iron manipulations are sensed by IRPs in the liver and spleen of wt and Hjv-/-mice

The IRE/IRP system orchestrates homeostatic adaptation to cellular iron supply (17, 18). To evaluate the responses of IRPs in the whole liver and spleen to the above-described dietary iron manipulations, we analyzed tissue extracts from wt and Hjv-/-mice by an electrophoretic mobility shift assay (EMSA) using a ^32^P-labelled IRE probe. The data in Fig. 5A-B show that HID intake tended to decrease the IRE-binding activities of IRP1 and IRP2 in both liver and spleen of wt mice (statistical significance is only reached in the liver); densitometric quantification of IRE/IRP1 and IRE/IRP2 complexes is shown on the right. Conversely, IDD intake significantly induced the IRE-binding activity of IRP2 in the liver and spleen of Hjv-/-mice, leaving IRP1 largely unaffected. IRE/IRP2 interactions are better visible in longer exposures (middle panels). EMSAs with tissue extracts previously treated with 2-mercaptoethanol (2-ME) were performed as loading controls (27) and are shown in the bottom panels.

**Fig. 5.**
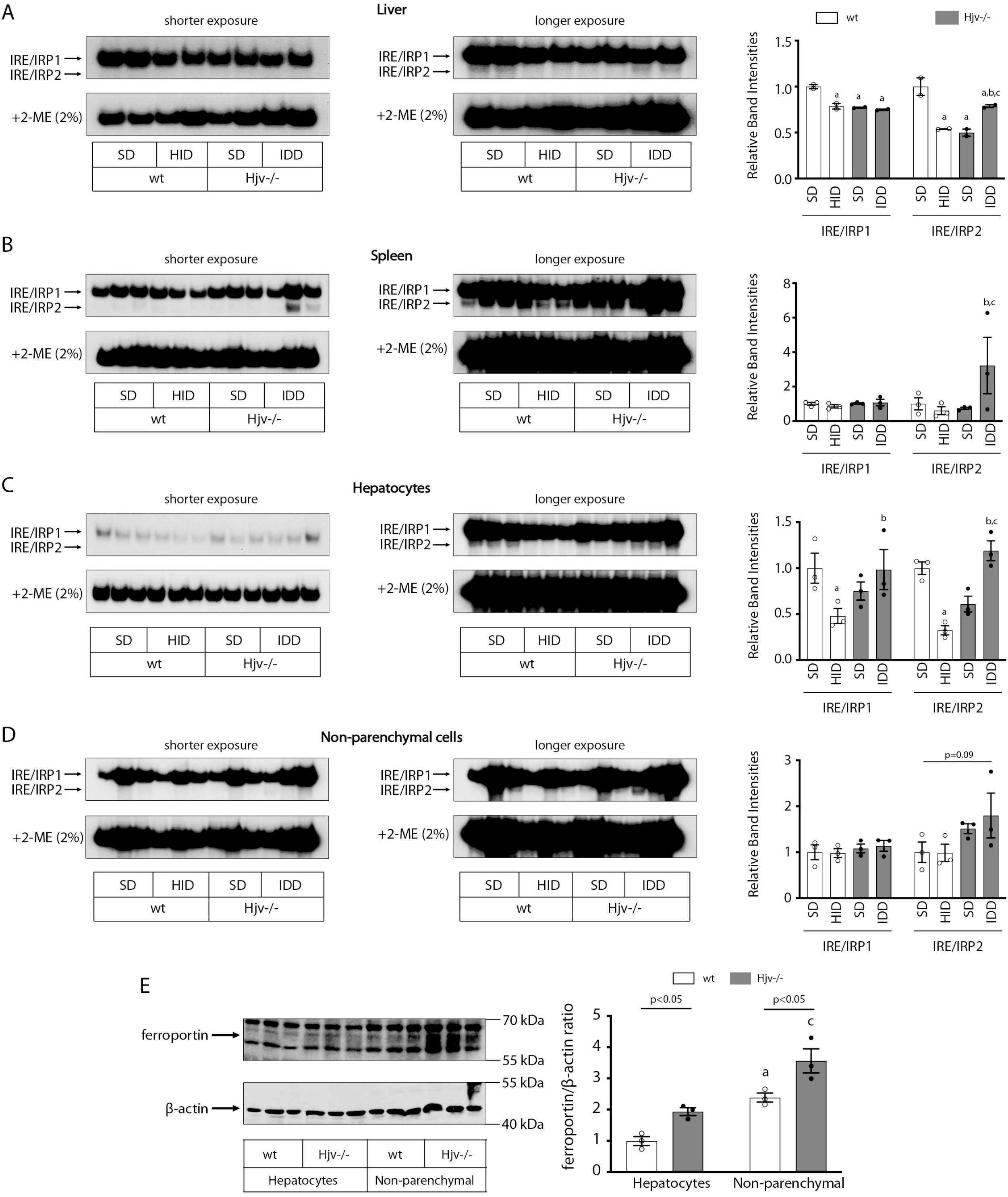
Dietary iron manipulations trigger IRP responses in the liver and spleen, as well as in primary hepatocytes and non-parenchymal liver cells of wt and Hjv-/-mice. Whole liver (A), whole spleen (B), isolated hepatocytes (C) or isolated non-parenchymal liver cells (D) from the mice described in Fig. 3 were analyzed for IRE-binding activity by EMSA with a ^32^P-labelled IRE probe in the absence (top) or presence (bottom) of 2% mercaptoethanol (2-ME). Two or three representative samples from each condition are shown. The positions of IRE/IRP1 and IRE/IRP2 complexes are indicated by arrows. Shorter and longer exposures of the autoradiograms are shown in the left and middle panels, respectively. Relative band intensities were quantified by densitometry and shown on the right panels. (E) Isolated hepatocytes and isolated non-parenchymal liver cells were analyzed by Western blotting for expression of ferroportin and β-actin. Blots were quantified by densitometry and ferroportin/β-actin ratios are shown on the right. Densitometric data are presented as the mean±SEM. Statistically significant differences (p<0.05) in values from control wt mice on SD are indicated by a, from wt mice on HID by b, and from Hjv-/-mice on SD by c.

To clarify which cell types of the liver account for the responses of IRPs to dietary iron, separate EMSAs were performed using extracts from isolated hepatocytes or non-parenchymal liver cells. The data in Fig. 5C-D uncover that IRP1 and IRP2 in both liver cell populations from wt and Hjv-/-mice are sensitive to dietary iron loading or restriction, respectively. The EMSA analysis of non-parenchymal liver cells, which contains Kupffer cells among others, showed a high experimental variability (Fig. 5D). Nevertheless, the overall results are consistent with those obtained with splenic extracts, which contain red pulp macrophages (Fig. 5B).

### Relative expression of ferroportin in hepatocytes and non-parenchymal liver cells from wt and Hjv-/-mice

We determined the relative abundance of ferroportin in hepatocytes and non-parenchymal liver cells from wt and Hjv-/-mice on SD by Western blotting. As expected, ferroportin expression (normalized to β-actin) was ∼1.5-2-fold higher in the non-parenchymal cell fraction as compared to hepatocytes in both wt and Hjv-/-mice (Fig. 5E). In comparison across genotypes, ferroportin expression was ∼2-fold higher in hepatocytes and ∼50% higher in non-parenchymal cells from Hjv-/-vs wt mice.

### Iron-dependent regulation of ferroportin mRNA translation in the liver

Having established that dietary iron manipulations trigger IRP responses in the liver and spleen, we hypothesized that the functional outcomes of exogenous hepcidin may not merely depend on its capacity to degrade tissue ferroportin, but also on iron-dependent ferroportin regeneration via *de novo* synthesis. *Fpn(+IRE)* mRNA is the predominant ferroportin transcript in the mouse liver and spleen, as well as in hepatoma and macrophage cell lines (21), and is considered as a target of IRPs.

Thus, we assessed the effects of dietary iron on whole liver *Fpn(+IRE)* mRNA translation by polysome profile analysis. We focused on the liver because this organ contains the highest number of iron-recycling macrophages (28) and can also export iron to plasma from ferroportin-expressing parenchymal cells. Liver extracts from wt mice on SD or HID, and Hjv-/-mice on SD or IDD were fractionated on sucrose gradients to separate translationally inactive light monosomes from translating heavy polysomes (Fig. 6A). The relative distribution of *Fpn(+IRE), Fth1* (positive control for iron regulation) and *Actb* (negative control) mRNAs within the different fractions was quantified by qPCR (Fig. 6B-D). Dietary iron loading stimulated *Fpn(+IRE)* (and *Fth1*) mRNA translation in wt mice (note the shifts from monosomes to polysomes in Fig. 6B-C). Conversely, dietary iron depletion inhibited *Fpn(+IRE)* (and *Fth1*) mRNA translation in Hjv-/-mice. We also attempted to obtain polysome profiles of *Fpn(-IRE)* mRNA but it was undetectable after fractionation. These data indicate that in mice subjected to iron overload, iron-stimulated ferroportin synthesis in the liver antagonizes hepcidin-mediated ferroportin degradation and prevents an appropriate hypoferremic response. Considering that levels of *Fpn(+IRE)* mRNA are elevated in iron-loaded wt and Hjv-/-mice (Figs. 1J and S7A), it is possible that increased *de novo* ferroportin synthesis is further enhanced by transcriptional induction.

**Fig. 6.**
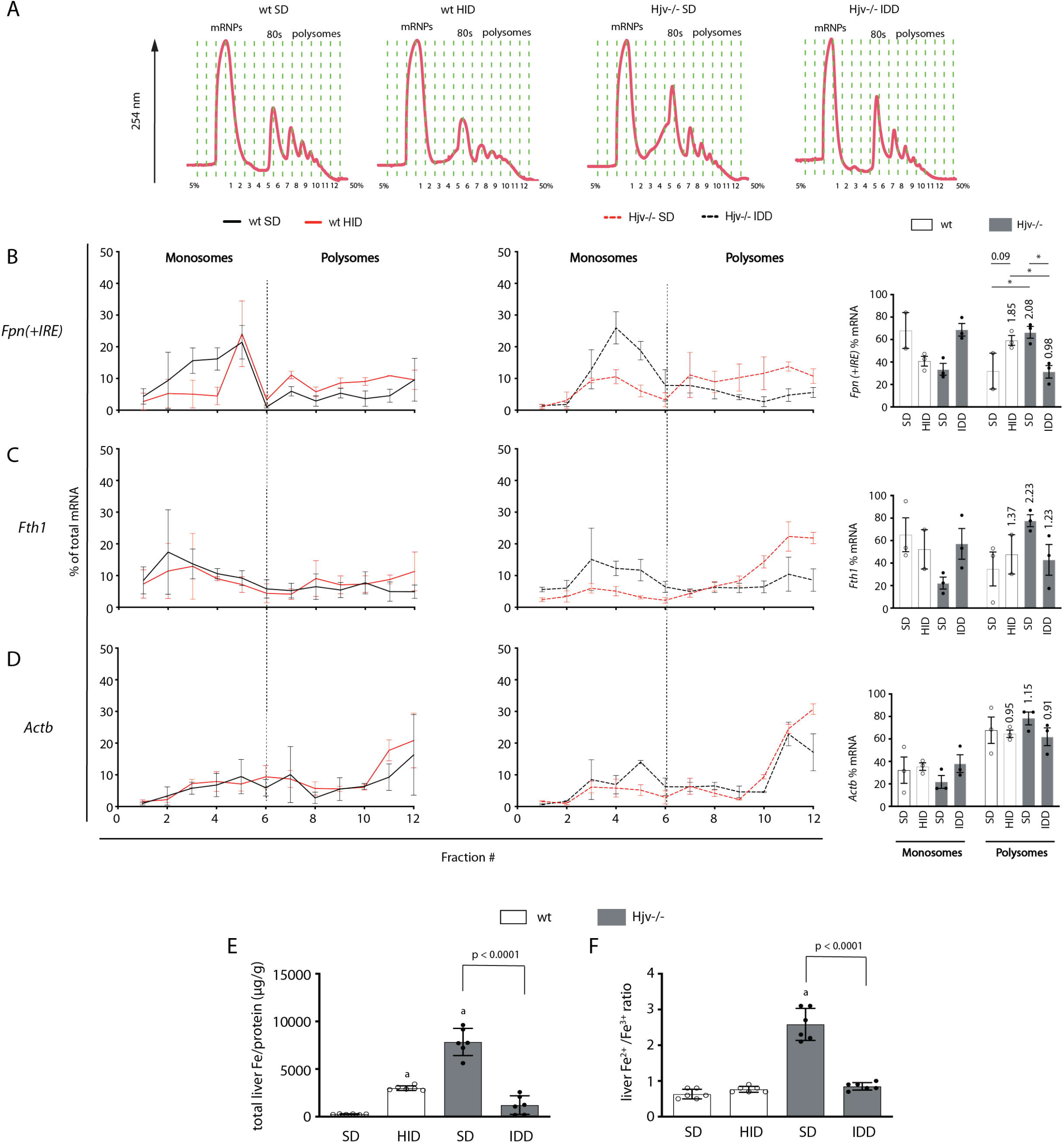
Iron regulation of Fpn(+IRE) mRNA translation in the mouse liver. Four-week-old wt male mice (n=10-14 per group) were placed on HID for five weeks. Conversely, age- and sex-matched isogenic Hjv-/-mice (n=10-14 per group) were placed on IDD for five weeks to prevent excessive iron overload. Controls from both genotypes were kept on SD. At the endpoint, the mice were sacrificed, and livers were used for polysome profile analysis and iron assays. (A) Recording of absorbance at 254 nm of representative samples. Fraction numbers and direction of the gradient are indicated. (B-D) Liver polysome profiles from n=3 mice in each experimental group. Distribution of (B) *Fpn(+IRE)*, (C) *Fth1* and (D) *Actb* mRNAs among light monosomal and heavy polysomal fractions (separated by dashed line) was analyzed by qPCR. Bar graph comparisons of pooled fractions are shown on the right. Numbers indicate the fold change compared to wt mice on SD. (E and F) Analysis of total iron (E), and redox iron speciation (F) in the liver by CE-ICP-MS. Data are presented as the mean±SEM. Statistical analysis in (A) was performed by two-way ANOVA and in (B, C) by one-way ANOVA. Statistically significant differences (p<0.05) compared to values from wt control mice on SD are indicated by a.

Quantification of liver iron by ICP-MS (Fig. 6E) validated iron loading of wt mice by HID, and relative iron depletion of Hjv-/-mice by IDD intake, respectively (see also Fig. 1D). Iron redox speciation analysis by CE-ICP-MS revealed a profound increase in Fe^2+^/Fe^3+^ ratios in livers of Hjv-/-mice on SD, which was corrected by dietary iron depletion (Fig. 6F). Nevertheless, there was no difference in Fe^2+^/Fe^3+^ ratios among livers of wt mice on SD or HID, and Hjv-/-mice on IDD. We conclude that a relative increase in total iron content, rather than excessive accumulation of redox active Fe^2+^ drives *Fpn(+IRE)* (and *Fth1*) mRNA translation in the liver.

### Restoration of effective hypoferremic response under iron overload following maximal Fpn mRNA suppression

We reasoned that complete inactivation of ferroportin mRNA would restore hepcidin-induced hypoferremia despite iron overload. An 8-hour treatment of mice with LPS suppressed liver *Fpn(+IRE)* mRNA below detection levels (Fig. 7A), as reported (9). The same holds true for the *Fpn(-IRE)* isoform (Fig. 7B), which was 290 times less abundant in control mouse livers compared to *Fpn(+IRE)* (ΔCt=8.18), in agreement with published data (21). We went on to examine the effects of synthetic hepcidin on serum iron under these conditions of maximal *Fpn* mRNA suppression. Importantly, the prolonged LPS treatment decreased serum iron in wt mice on HID below the control baseline (Fig. 7C). Furthermore, when combined with synthetic hepcidin, it promoted an effective hypoferremic response in wt mice on HID and Hjv-/-mice on SD (or IDD) (Figs. 7C-D) and tended to decrease NTBI (Fig. 7E). These data strongly suggest that the expression of actively translating *Fpn* mRNA in iron-exporting tissues under systemic iron overload mitigates hepcidin-induced drop in serum iron.

**Fig. 7.**
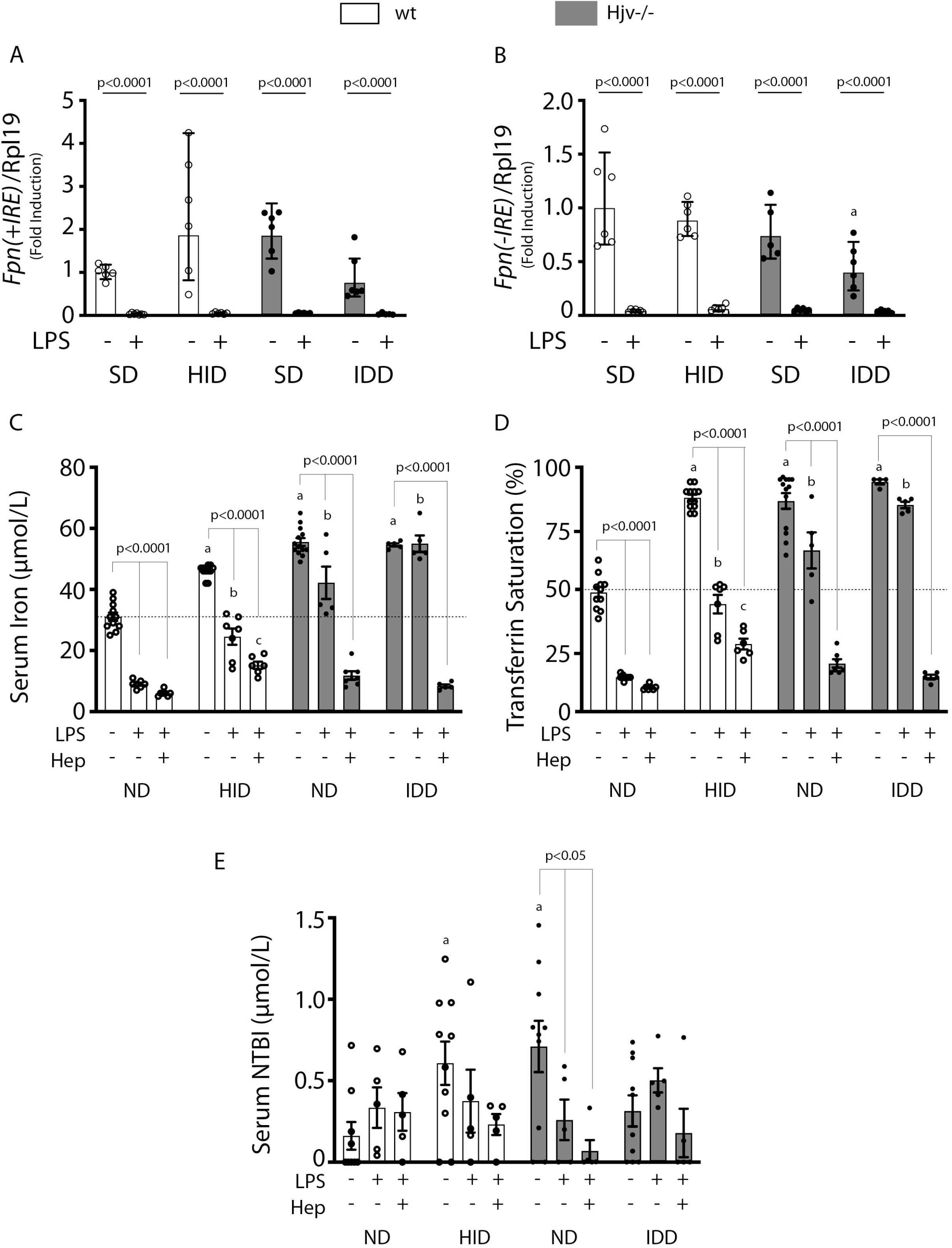
Elimination of ferroportin mRNA by prolonged LPS treatment potentiates hepcidin-induced hypoferremia in mouse models of iron overload. Four-week-old wt male mice (n=10-14 per group) were placed on HID for five weeks. Conversely, age- and sex-matched isogenic Hjv-/-mice (n=10-14 per group) were placed on IDD for five weeks to prevent excessive iron overload. Controls from both genotypes were kept on SD. (A and B) Half of the mice were injected with saline and the other half with 1 µg/g LPS and sacrificed after 8 h. Livers were dissected and processed for qPCR analysis of *Fpn(+IRE)* (A) and *Fpn(-IRE)* (B) mRNAs. (C-E) All mice were injected with 1 µg/g LPS. Half of the animals were subsequently injected with saline, and the other half with 2.5 µg/g synthetic hepcidin every two hours for a total of 4 injections. At the endpoint the mice were sacrificed. Sera were collected by cardiac puncture and analyzed for: (C) iron, (D) transferrin saturation, and (E) NTBI. The dotted line in (C) and (D) indicates baseline serum iron and transferrin saturation, respectively, of control wt mice on SD. Data are presented as (A-B) geometric mean±SD or (C-E) mean±SEM. Statistically significant differences (p<0.05) compared to values from saline-, LPS- or hepcidin-treated wt control mice on SD are indicated by a, b or c, respectively.

## Discussion

We sought to analyze how iron overload affects hepcidin-mediated inflammatory responses. We and others reported that excess iron inhibits the major hepcidin signaling pathways (BMP/SMAD and IL-6/STAT3) in cultured cells (10, 11). To explore the physiological relevance of these findings, wt mice were subjected to variable degrees of dietary iron loading and then treated with LPS. All iron-loaded mice could further upregulate hepcidin in response to LPS-induced acute inflammation (Fig. S1). This is consistent with other relevant findings (29) and apparently contradicts the *in vitro* data. While experimental iron loading of cultured cells is rapid, dietary iron loading of mice is gradual (20) and most of excess iron is effectively detoxified within ferritin, which is highly induced (30). By contrast, the suppression of hepcidin preceded ferritin induction in cultured cells (10), which may explain the discrepancy with the *in vivo* data.

The unimpaired inflammatory induction of hepcidin in iron-loaded wt mice correlated with significant drops in serum iron, but these appeared inversely proportional to the degree of systemic iron loading (Fig. S1). Thus, LPS-treated mice on 5 weeks HID developed relative hypoferremia but could not further reduce serum iron below a baseline of untreated control mice on SD. This can be attributed to mechanisms antagonizing hepcidin action. To explore how iron modulates the capacity of hepcidin to trigger inflammatory hypoferremia, we established conditions of iron overload using wt and Hjv-/-mice with extreme differences in hepcidin expression. Figs. 1 and 2 demonstrate that iron overload prevents effective inflammatory hypoferremia independently of hepcidin and tissue ferroportin levels.

We used a ∼200-fold excess of synthetic hepcidin to directly assess its capacity to divert iron traffic in iron-loaded mice. Hepcidin injection caused hypoferremia in control wt mice on SD and significantly reduced serum iron in wt mice on HID and Hjv-/-mice on SD, but not below baseline (Fig. 3). Thus, synthetic hepcidin failed to drastically drop serum iron levels in iron overload models with either high or low endogenous hepcidin. Importantly, synthetic hepcidin promoted robust hypoferremia in relatively iron-depleted Hjv-/-mice on IDD, with undetectable endogenous hepcidin. It should be noted that synthetic hepcidin had similar effects on tissue ferroportin among wt or Hjv-/-mice, regardless of iron diet (Fig. 4). It reduced intensity of the ferroportin signal in Kupffer cells and splenic macrophages of wt mice without significantly affecting biochemically detectable total protein levels. In addition, it dramatically reduced total ferroportin in the liver and spleen of Hjv-/-mice. However, in all experimental settings there was residual tissue ferroportin, which appears to be functionally significant.

We reasoned that at steady-state, tissue ferroportin may consist of fractions of newly synthesized protein, and protein that is *en route* to hepcidin-mediated degradation. Conceivably, the former may exhibit more robust iron export activity, at least before its iron channel gets occluded by hepcidin. Increased *de novo* synthesis of active ferroportin could explain why synthetic hepcidin cannot drastically drop serum iron levels under iron overload. In fact, Fig. 6 demonstrates that dietary iron overload augments *Fpn(+IRE)* mRNA translation in the liver of wt mice. Conversely, relative dietary iron depletion inhibits *Fpn(+IRE)* mRNA translation in the liver of Hjv-/-mice, in line with the restoration of hepcidin-mediated hypoferremic response (Fig. 3).

Our data are consistent with translational control of liver ferroportin expression via the IRE/IRP system and do not exclude the possibility for an additional contribution of iron-dependent transcriptional regulation of *Fpn(+IRE)* mRNA. Direct evidence for activation of IRP responses in the liver and spleen to dietary iron manipulations is provided in Fig. 5. While translational control of ferritin in tissues is established (31), regulation of ferroportin by the IRE/IRP system is less well characterized and has hitherto only been documented in cell models (32, 33), the mouse duodenum (34), and the rat liver (35). Moreover, the physiological relevance of this mechanism remained speculative. The data in Figs. 5 and 6 show that the IRE/IRP system is operational and controls *Fpn(+IRE)* mRNA translation in both fractions of hepatocytes and non-parenchymal liver cells. Presumably, this offers a compensatory mechanism to protect the cells from iron overload and iron-induced toxicity. On the other hand, this mechanism attenuates hepcidin responsiveness and promotes a state of hepcidin resistance, as higher amounts of hepcidin are required to achieve effective hypoferremia. Because hepcidin has a short plasma half-life, it is reasonable to predict that the use of more potent hepcidin analogs (36) will overcome the antagonistic effects of increased ferroportin mRNA translation under iron overload.

The critical role of *de novo* ferroportin synthesis in fine-tuning hepcidin-dependent functional outcomes is also highlighted in Fig. 7. Thus, synthetic hepcidin was highly effective as promoter of hypoferremia in dietary iron-loaded wt mice when administered together with LPS. LPS is known to suppress *Fpn* mRNA in cell lines (16) and mouse tissues, with a nadir in the liver reached at 8 h (9). The recovery of hepcidin effectiveness in mouse models of iron overload was only possible when *Fpn* mRNA was essentially eliminated. Under these conditions, LPS treatment alone was sufficient to decrease serum iron in dietary iron-loaded wt mice below baseline.

Tissue iron uptake may be another important determinant of the hypoferremic response to inflammation. LPS did not affect *Tfrc* mRNA levels in the liver (Fig. 1O), which argues against increased uptake of transferrin-bound iron via TfR1. On the other hand, LPS induced *Zip14, Dmt1* and *Lcn2* mRNAs (Figs 1L-N). Zip14 is the NTBI transporter accounting for hepatocellular iron overload in hemochromatosis (37) and is upregulated by inflammatory cues in hepatocytes (38). DMT1 is dispensable for NTBI uptake by hepatocytes (39), and its inflammatory induction might promote iron acquisition by macrophages (16, 40). Nevertheless, considering that the fraction of NTBI represents <2% of total serum iron even in the iron overload models (Figs. 1A and C), it is implausible that NTBI uptake by Zip14 and/or DMT1 substantially contributes to inflammatory hypoferremia. Lcn2 is an acute phase protein that can sequester intracellular iron bound to catecholate siderophores (41), and is more likely to transport iron to tissues during infection. In any case, synthetic hepcidin did not affect expression of iron transporters (Fig. S7E-H). This excludes the possibility for a synergistic effect on LPS-induced tissue iron uptake that could promote effective hypoferremia in the iron overload models.

Our study has some limitations. While the data highlight the importance of translational regulation of liver ferroportin as a determinant of serum iron, they do not accurately dissect the specific role of ferroportin expressed in hepatocytes and Kupffer cells; the latter were not separated from other non-parenchymal cells in biochemical assays. The involvement of the IRE/IRP system has been established indirectly, while the relative contributions of IRP1 and IRP2 in the mechanism are not fully defined. The possible role of iron-dependent transcriptional induction of ferroportin in counterbalancing hepcidin actions requires further clarification. The use of diets with variable iron content may have triggered responses to iron availability independent of hepcidin signaling and Hjv functionality. Finally, the physiological implications of translational regulation of ferroportin in the broader setting of inflammation and/or infection have not been explored.

In conclusion, our data reveal a crosstalk between the hepcidin pathway and the IRE/IRP system in the liver and spleen for the control of tissue ferroportin and serum iron levels. Furthermore, they suggest that application of hepcidin therapeutics for treatment of iron overload disorders should be combined with iron depletion strategies to mitigate *Fpn* mRNA translation and increase hepcidin efficacy. Future work is expected to clarify whether optimizing the hypoferremic response to inflammation under systemic iron overload decreases susceptibility to pathogens.

## Materials and Methods

### Animals

Wild type C57BL/6J and isogenic Hjv-/-mice (42) were housed in macrolone cages (up to 5 mice/cage, 12:12 h light-dark cycle: 7 am - 7 pm; 22 ± 1°C, 60 ± 5% humidity). The mice were fed either a standard diet (200 ppm iron), an iron-deficient diet (2-6 ppm iron) or a high-iron diet (2% carbonyl iron) (43). Where indicated, mice were injected intraperitoneally with 1 μg/g LPS (serotype 055:B5; Sigma-Aldrich) or subcutaneously with 2.5 µg/g synthetic hepcidin; control mice were injected with phosphate-buffered saline. At the endpoints, animals were sacrificed by CO_2_ inhalation and cervical dislocation. Experimental procedures were approved by the Animal Care Committee of McGill University (protocol 4966).

### Serum biochemistry

Blood was collected via cardiac puncture. Serum was prepared by using micro Z-gel tubes with clotting activator (Sarstedt) and was kept frozen at −20°C until analysis. Serum iron, total iron binding capacity (TIBC) and, where indicated serum ferritin, were determined at the Biochemistry Department of the Montreal Jewish General Hospital using a Roche Hitachi 917 Chemistry Analyzer. Transferrin saturation was calculated from the ratio of serum iron and TIBC. Serum hepcidin was measured by using an ELISA kit (HMC-001; Intrinsic LifeSciences).

### Quantification of serum non-transferrin bound iron (NTBI)

NTBI was measured by adapting the method developed by Esposito *et al* (44). Briefly, iron samples of known concentration were created by mixing 70 mM nitrilotriacetate (NTA) (pH = 7.0) with 20 mM ferrous ammonium sulfate. Fe^2+^ was allowed to oxidize to Fe^3+^ in ambient air for at least 30 min and then the solution was diluted to 0.2 mM before further serial dilutions to create a ladder. 5 μl of ladder was loaded in a 96-well plate containing 195 μl plasma-like medium with or without 100 μM deferiprone. The composition of the plasma-like medium was: 40 mg/ml bovine serum albumin, 1.2 mM sodium phosphate dibasic, 120 μM sodium citrate, 10 mM sodium bicarbonate in iron-free HEPES-buffered saline (HEPES 20 mM, NaCl 150 mM, treated with Chelex-100 chelating resin [Bio-Rad, Hercules, CA], 0.5 mM NTA, 40 μM ascorbic acid, 50 μM dihydrorhodamine, pH=7.4). 5 μl of sample was loaded in a 96-well plate containing 195 μl of iron-free HEPES-buffered saline with or without 100 μM deferiprone. Microplates were read every 2 minutes at 37□ over 40 min at 485/520 nm (ex/em). Final NTBI was calculated by comparing the oxidation rate of DHR in the presence or absence of the strong chelator deferiprone.

### Hepcidin synthesis

Human hepcidin (DTHFPICIFCCGCCHRSKCGMCCKT) was synthesized at Ferring Research Institute, San Diego, CA. The linear peptide was assembled on Rink amide resin using Tribute peptide synthesizer and the peptide was cleaved from the resin with the TFA/TIS/EDT/H_2_O 91:3:3:3 (v/v/v/v) cocktail. The solvents were evaporated, and the crude peptide was precipitated with diethyl ether, reconstituted in 50% aqueous acetonitrile and lyophilized. The lyophilizate was dissolved in 30% aqueous acetonitrile at the concentration of 0.05 mM and the pH of the solution was adjusted to 7.8 with 6 M ammonium hydroxide. Folding was achieved within 4 hours using the cysteine/cystine redox (peptide/Cys/Cys_2_ 1:6:6 molar ratio). The reaction mixture was acidified to pH 3, loaded onto HPLC prep column and purified in a TFA based gradient. The identity of the peptide was confirmed by mass spectrometry and by coelution with a commercially available sample (Peptide International, #PLP-3771-PI).

### Quantitative real-time PCR (qPCR)

RNA was extracted from livers by using the RNeasy kit (Qiagen). cDNA was synthesized from 1 μg RNA by using the OneScript® Plus cDNA Synthesis Kit (Applied Biological Materials Inc.). Gene-specific primers pairs (Table S1) were validated by dissociation curve analysis and demonstrated amplification efficiency between 90-110 %. SYBR Green (Bioline) and primers were used to amplify products under following cycling conditions: initial denaturation 95°C 10 min, 40 cycles of 95°C 5 s, 58°C 30 s, 72°C 10 s, and final cycle melt analysis between 58°-95°C. Relative mRNA expression was calculated by the 2^-ΔΔ^Ct method (45). Data were normalized to murine ribosomal protein L19 (*Rpl19*). Data are reported as fold increases compared to samples from wild type mice on standard diet (SD).

### Polysome fractionation

RNA was freshly prepared from frozen livers. Linear sucrose gradients were prepared the day before the experiment by using 5% (w/v) and 50% (w/v) sucrose solutions with 10x gradient buffer (200 mM HEPES pH=7.6, 1 M KCl, 50 mM MgCl_2_, 0.1 mg/ml Cycloheximide, 1 tablet cOmplete™, Mini, EDTA-free Protease Inhibitor Cocktail (Roche), 200 U/mL Recombinant RNasin® Ribonuclease Inhibitor (Promega), 2 mM DTT). Linear gradients were prepared in Polyallomer Centrifuge Tubes (Beckman Coulter). Tubes were marked using a gradient cylinder (BioComp), and 5% sucrose solution was added using a syringe with a layering needle (BioComp) until solution level reached the mark. Then, 50% sucrose solution was layered underneath the 5% solution until the interface between the two solutions reached the mark. Tubes were capped with rate zonal caps (BioComp) and linearized using a Gradient Master 108 (Biocomp). All reagents were nuclease-free and all solutions were kept on ice or at 4□. Sample preparation was adapted from Liang *et al*. (46). Briefly, livers were flash frozen upon collection. Roughly 30-80 mg of tissue was crushed using a mortar and pestle in the presence of liquid nitrogen to prevent thawing. Tissues were lysed in up to 1 ml of hypotonic lysis buffer (5 mM Tris-Hcl pH=7.5, 1.5 mM KCl, 2.5 mM MgCl_2_, 2 mM DTT, 1 mg/ml Cycloheximide, 200 U/ml Recombinant RNasin® Ribonuclease Inhibitor (Promega), 1 tablet cOmplete™, Mini, EDTA-free Protease Inhibitor Cocktail (Roche), 0.5% (v/v) Triton X-100, 0.5% (v/v) Sodium Deoxycholate) and homogenized using Dounce homogenizers (60 movements with both loose and tight pestles) on ice. Samples were centrifuged at 4□, 16,060g for 4 minutes and supernatants were collected. Sample optical density was measured at 260 nM and samples were normalized to either the lowest value or 30 ODs. 450 μl of sucrose gradient was removed from the top and replaced with normalized sample. Tube weights were balanced by weight before centrifugation at 200,000g for 2 h at 4□ in a SW 41 Ti rotor and a Beckman Optima L-60 Ultracentrifuge. Samples were fractionated using a BR-188 Density Gradient Fractionation System (Brandel). Immediately upon collection, 800 μl of samples were mixed with 1 ml of TRIzol™ and kept on ice before storage at -80□. Polysomal RNA was processed according to the manufacturer’s protocol. mRNA distribution was analyzed as previously described (47).

### Electrophoretic mobility shift assay (EMSA)

IRE-binding activities from liver and spleen were analyzed by EMSA using a radioactive ^32^P-labelled IRE probe, according to established procedures (27). EMSAs were also performed in extracts from hepatocytes and non-parenchymal cells, which were separated by using a 2-step collagenase perfusion technique, as previously described (9).

### Western blotting

Livers were washed with ice-cold PBS and dissected into pieces. Aliquots were snap frozen at liquid nitrogen and stored at −80°C. Protein lysates were obtained as described (22). Lysates containing 40 μg of proteins were analyzed by SDS-PAGE on 9-13% gels and proteins were transferred onto nitrocellulose membranes (BioRad). The blots were blocked in non-fat milk diluted in tris-buffered saline (TBS) containing 0.1% (v/v) Tween-20 (TBS-T), and probed overnight with antibodies against ferroportin (48) (1:1000 diluted monoclonal rat anti-mouse 1C7, kindly provided by Amgen Inc), β-actin (1:2000 diluted; Sigma), or Tfr2 (1:1000 diluted rabbit polyclonal; Alpha Diagnostics). Following a 3x wash with TBS-T, the membranes were incubated with peroxidase-coupled secondary antibodies for 1 h. Immunoreactive bands were detected by enhanced chemiluminescence with the Western Lightning ECL Kit (Perkin Elmer).

### Immunohistochemistry

Tissue specimens were fixed in 10% buffered formalin and embedded in paraffin. Samples from 3 different mice for each experimental condition were cut at 4-µm, placed on SuperFrost/Plus slides (Fisher) and dried overnight at 37°C. The slides were then loaded onto the Discovery XT Autostainer (Ventana Medical System) for automated immunohistochemistry. Slides underwent de-paraffinization and heat-induced epitope retrieval. Immunostaining was performed by using 1:500 diluted rabbit polyclonal antibodies against ferroportin (49) and an appropriate detection kit (Omnimap rabbit polyclonal HRP, #760-4311 and ChromoMap-DAB #760-159; Roche). Negative controls were performed by the omission of the primary antibody. Slides were counterstained with hematoxylin for four minutes, blued with Bluing Reagent for four minutes, removed from the autostainer, washed in warm soapy water, dehydrated through graded alcohols, cleared in xylene, and mounted with Permount (Fisher). Sections were analyzed by conventional light microscopy and quantified by using the Aperio ImageScope software (Leica Biosystems) (9).

### Perls Prussian blue staining

To visualize non-heme iron deposits, deparaffinized tissue sections were stained with Perls’ Prussian blue using the Accustain Iron Stain kit (Sigma).

### Quantification of liver iron content (LIC)

Total liver iron was quantified by using the ferrozine assay (20) or inductively coupled plasma mass spectrometry (ICP-MS) (50).

### Iron speciation analysis

Iron redox speciation analysis in the liver was performed by capillary electrophoresis (CE) coupled to ICP-MS (CE-ICP-MS). Dynamic reaction cell (DRC) technology (ICP-DRC-MS) with NH_3_ as DRC-gas was applied for non-interfered monitoring of the iron isotopes. A “PrinCe 706” CE system (PrinCe Technologies B.V., Emmen, Netherlands) was employed for separation of iron species at +20 kV. Temperature settings for sample/buffer tray and capillary were set to 20°C. An uncoated capillary (100 cm x 50 µm ID; CS-Chromatographie Service GmbH, Langerwehe, Germany) was used for separation and hyphenation to the ICP–DRC-MS. A CE-ICP-MS interface (50, 51) was installed which provided the electrical connection between CE capillary end and outlet electrode. The self-aspiration mode allowed for best flow rate adjustment and avoided suction flow. Electrolytes for sample stacking and electrophoretic separation were 10% HCl = leading electrolyte, 0.05 mM HCl= terminating electrolyte and 50 mM HCl = background electrolyte. The Fe^2+^/Fe^3+^ ratio was calculated from quantitative determined concentrations of Fe-species.

### Statistics

Statistical analysis was performed by using the Prism GraphPad software (version 9.1.0). Lognormally distributed data including qPCR and ELISA results were first log transformed before analysis with ordinary two-way ANOVA (Tukey’s multiple comparisons test) for comparisons within same treatment groups (denoted by a or b on figures) or with multiple unpaired t tests using the Holm-Sidak method to compare effects between treatments. Normally distributed data was analyzed by two-way ANOVA using either Sidak’s method for comparisons between treatment groups or Tukey’s multiple comparisons test within treatments groups. Where indicated, pairwise comparisons were done with unpaired Student’s t test. Probability value p<0.05 was considered statistically significant.

## Acknowledgments

We thank Dr. Naciba Benlimame and Lilian Canetti for assistance with histology and immunohistochemistry. This work was supported by a grant from the Canadian Institutes of Health Research (CIHR; PJT-159730). EC was funded by a fellowship from the Natural Sciences and Engineering Research Council of Canada (NSERC) and is currently a recipient of a fellowship from the *Fonds de recherche du Québec – Santé* (FRQS). The work of VV and BM was financially supported by the Deutsche Forschungsgemeinschaft (DFG) through the Priority Program “Ferroptosis: from Molecular Basics to Clinical Applications” (SPP 2306).

## Figures legends

**Table S1.**
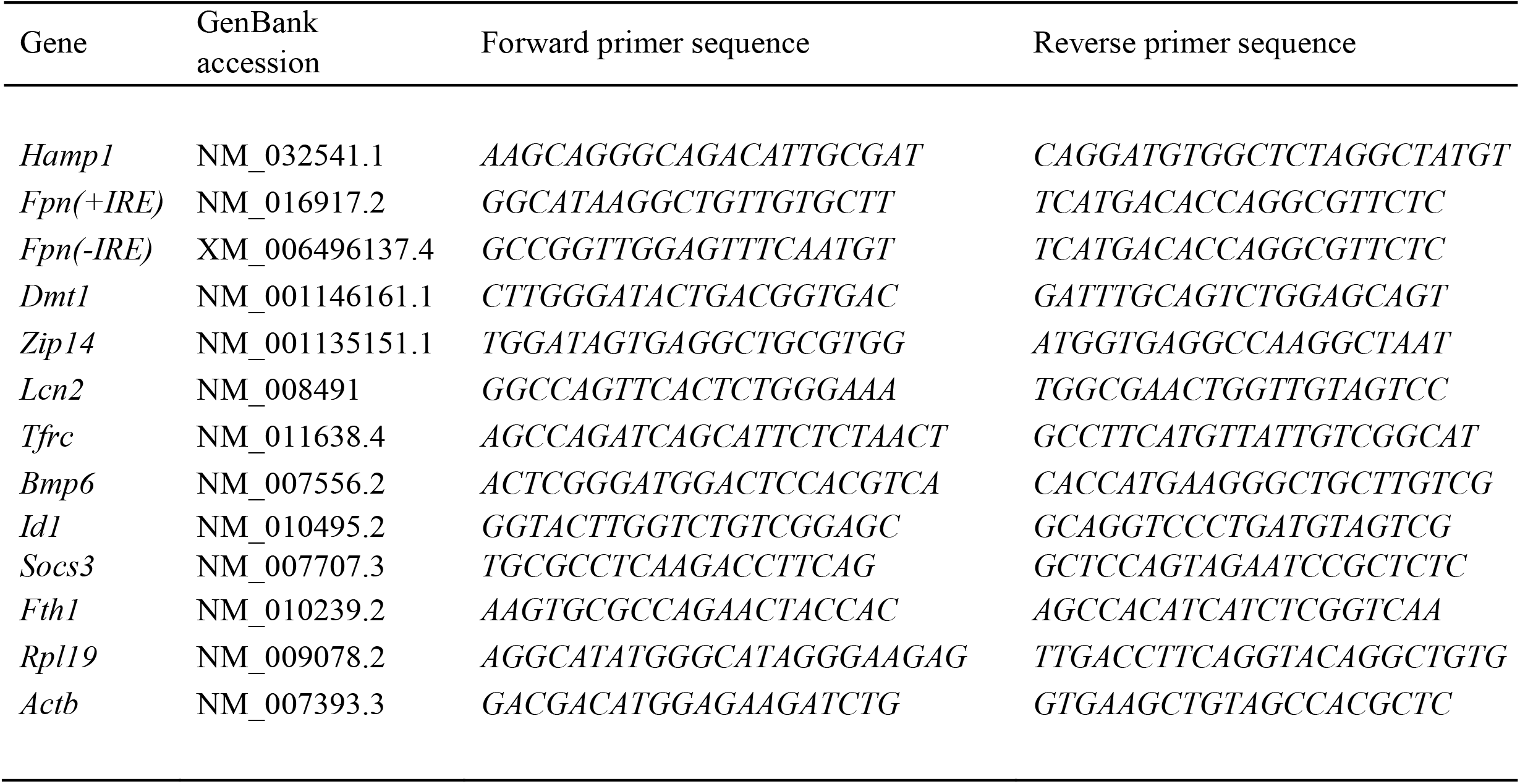
List of primers used for qPCR.

## Supplemental Figure legends

**Fig. S1. Dietary iron loading does not disrupt inflammatory hepcidin induction in LPS-treated wild type mice but blunts hepcidin-mediated hypoferremia**. Nine-week-old male mice (n=12-14 per group) were fed SD or HID for one day, one week, or five weeks prior to sacrifice. Half of the mice were injected intraperitoneally with saline and the other half with 1 µg/g LPS 4 hours before sacrifice. Sera were collected by cardiac puncture and analyzed for: (A) iron, (B) transferrin saturation, (D) ferritin, and (E) hepcidin. Livers were dissected and processed for biochemical analysis of: (C) liver iron content (LIC) by the ferrozine assay and (F) *Hamp* mRNA by qPCR. The dotted line in (A) and (B) indicates baseline serum iron and transferrin saturation, respectively, of control mice on SD. Data (A-E) are presented as the mean±SEM and in (F) as geometric mean±SD. Statistically significant differences (p<0.05) over time compared to values from saline- or LPS-treated control mice are indicated by a or b, respectively.

**Fig. S2. Effects of dietary iron manipulations in hepatic and splenic iron of wt and Hjv-/-mice**. (A) Liver and (B) spleen sections from mice described in Fig. 1 were analyzed histologically for iron deposits by Perls staining (magnification: 10x).

**Fig. S3. Low magnification immunohistochemical images of ferroportin in liver sections of dietary iron-manipulated wild type and Hjv-/-mice following LPS treatment**. Liver sections from mice described in Fig. 1 were used for immunohistochemical analysis of ferroportin (magnifications: 10x and 5x).

**Fig. S4. Low magnification immunohistochemical images of ferroportin in spleen sections of dietary iron-manipulated wild type and Hjv-/-mice following LPS treatment**. Spleen sections from mice described in Fig. 1 were used for immunohistochemical analysis of ferroportin (magnification: 2x).

**Fig. S5. Perls staining for iron deposits in liver and spleen sections of dietary iron-manipulated wild type and Hjv-/-mice following treatment with synthetic hepcidin**. Liver and spleen sections from mice described in Fig. 3 were stained with Perls Prussian blue (magnification: 10x).

**Fig. S6. Western analysis of transferrin receptors of dietary iron-manipulated wild type and Hjv-/-mice following treatment with synthetic hepcidin**. Livers from mice described in Fig. 3 were analyzed by Western blot for expression of Tfrc, Tfr2, and β-actin; a representative image (out of n=2 samples) is shown on the left. The blots were quantified by densitometry and Tfrc/β-actin or Tfr2/β-actin ratios are shown on the right.

**Fig. S7. Effects of LPS treatment on expression of mRNAs encoding iron transport proteins and signaling endpoints in the liver of dietary iron-manipulated wild type and Hjv-/-mice**. Livers from mice described in Fig. 3 were dissected and processed for qPCR analysis of mRNAs encoding iron transport proteins and signaling endpoints. (A) *Fpn(+IRE)*, (B) *Socs3*, (C) *Id1*, (D) *Bmp6*, (E) *Dmt1*, (F) *Zip14*, (G) *Lcn2* and (H) *Tfrc*. All data are presented as the geometric mean

± SD. Statistically significant differences (p<0.05) compared to values from saline- or hepcidin-treated control mice are indicated by a or b, respectively.

**Fig. S8. Low magnification immunohistochemical images of ferroportin in liver sections of dietary iron-manipulated wild type and Hjv-/-mice following treatment with synthetic hepcidin**. Liver sections from mice described in Fig. 3 were used for immunohistochemical analysis of ferroportin (magnifications: 10x and 5x).

**Fig. S9. Low magnification immunohistochemical images of ferroportin in spleen sections of dietary iron-manipulated wild type and Hjv-/-mice following treatment with synthetic hepcidin**. Spleen sections from mice described in Fig. 3 were used for immunohistochemical analysis of ferroportin (magnification: 2x).

