## Supplementary figures and images for "A crosstalk between hepcidin and IRE/IRP pathways controls ferroportin expression and determines serum iron levels in mice"

### Supplemental Figures

#### Slide 1
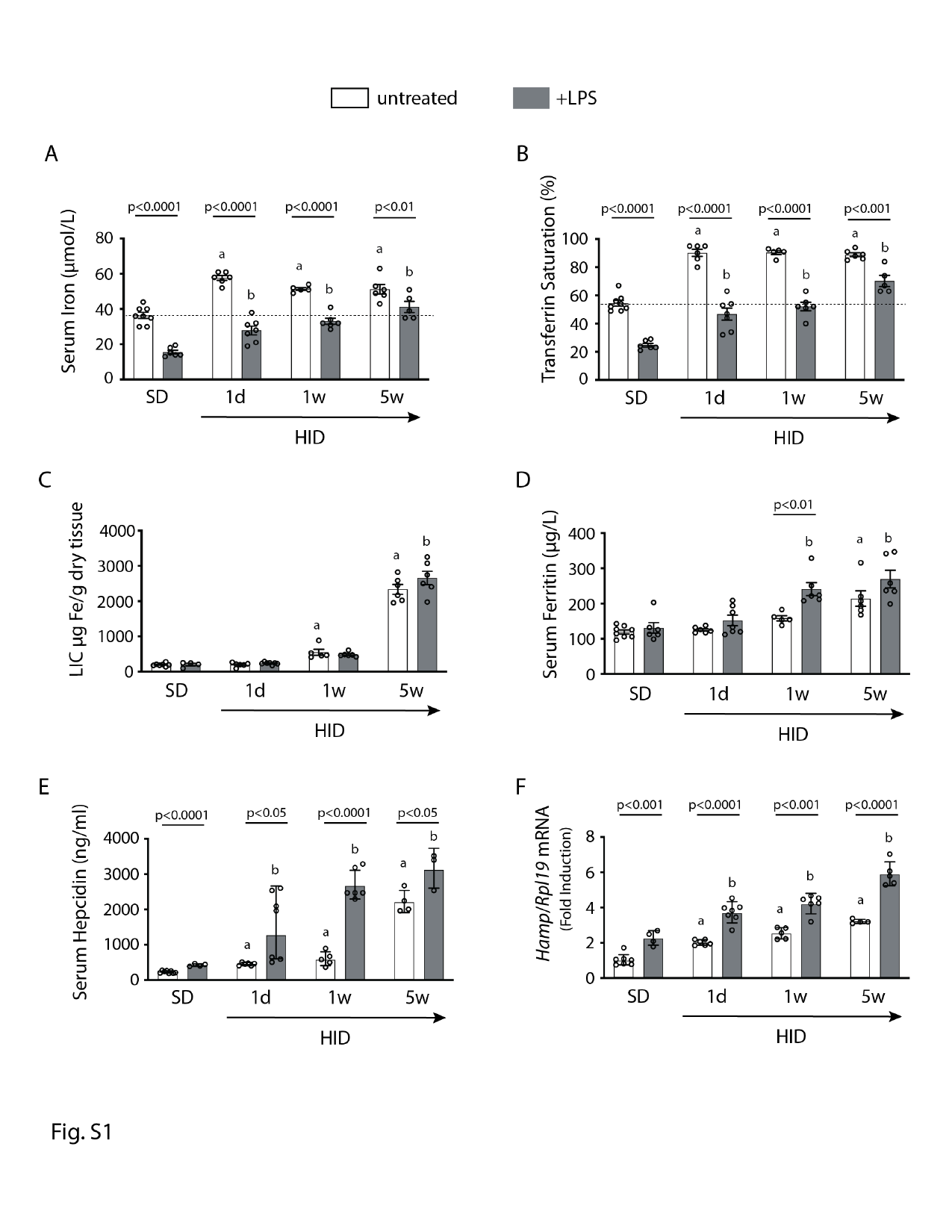

#### Slide 2
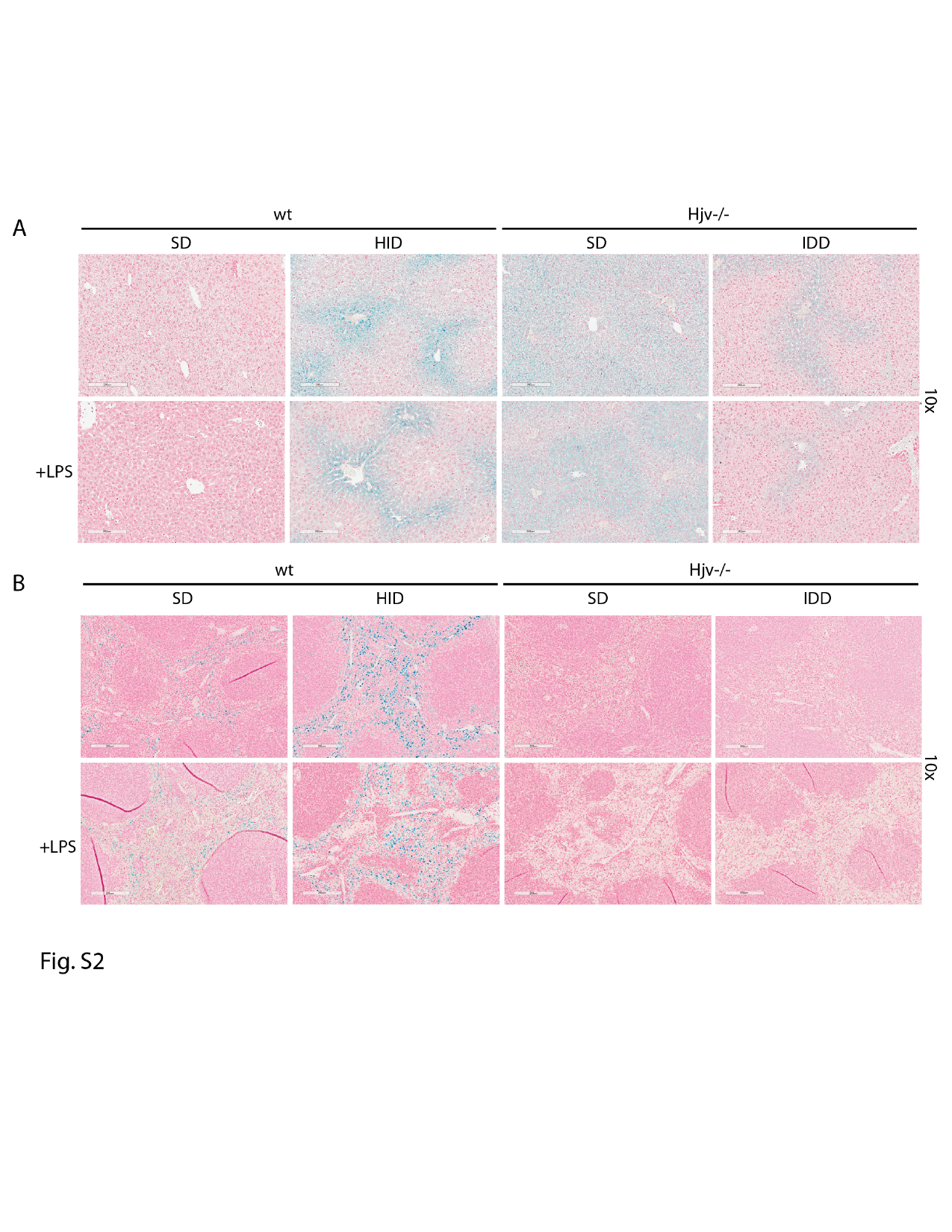

#### Slide 3
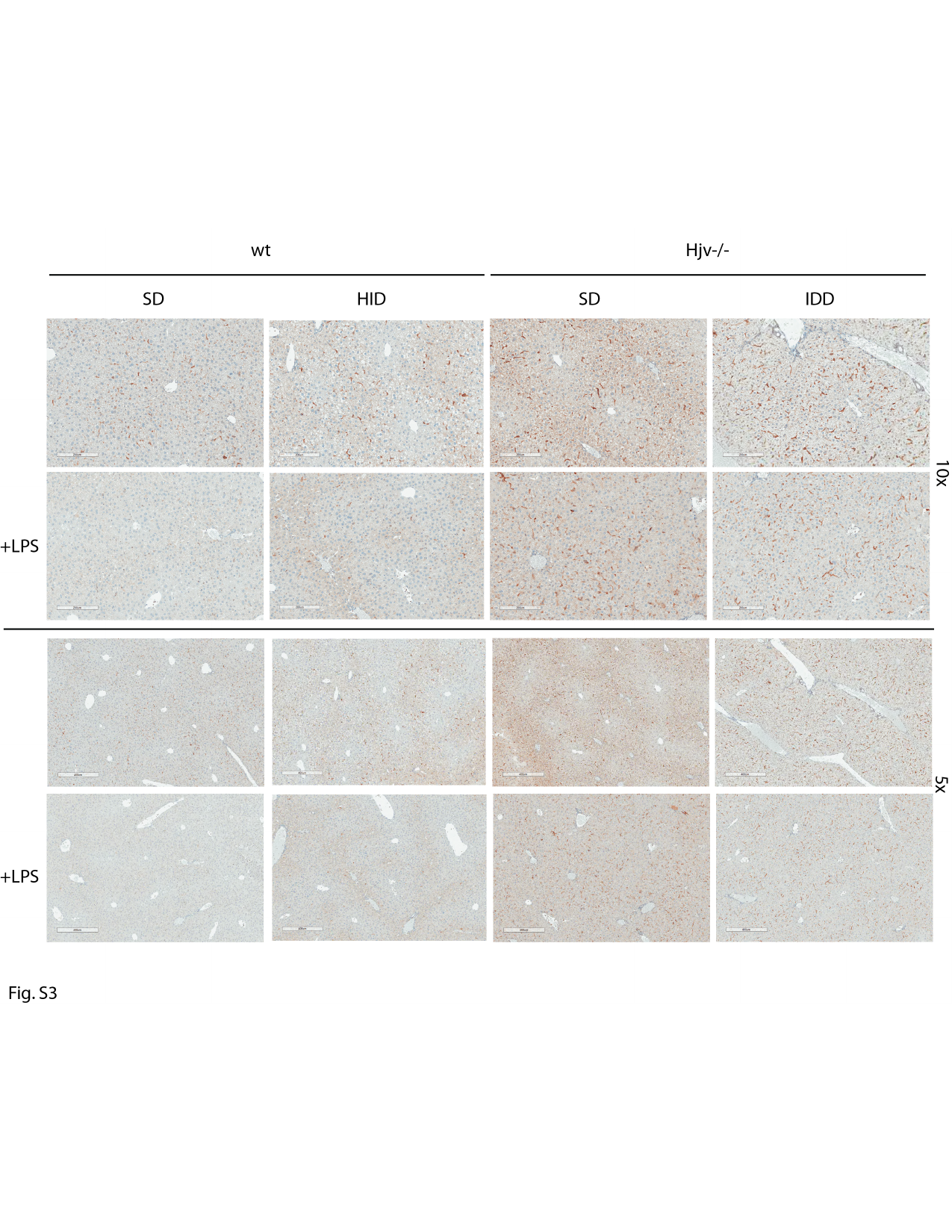

#### Slide 4
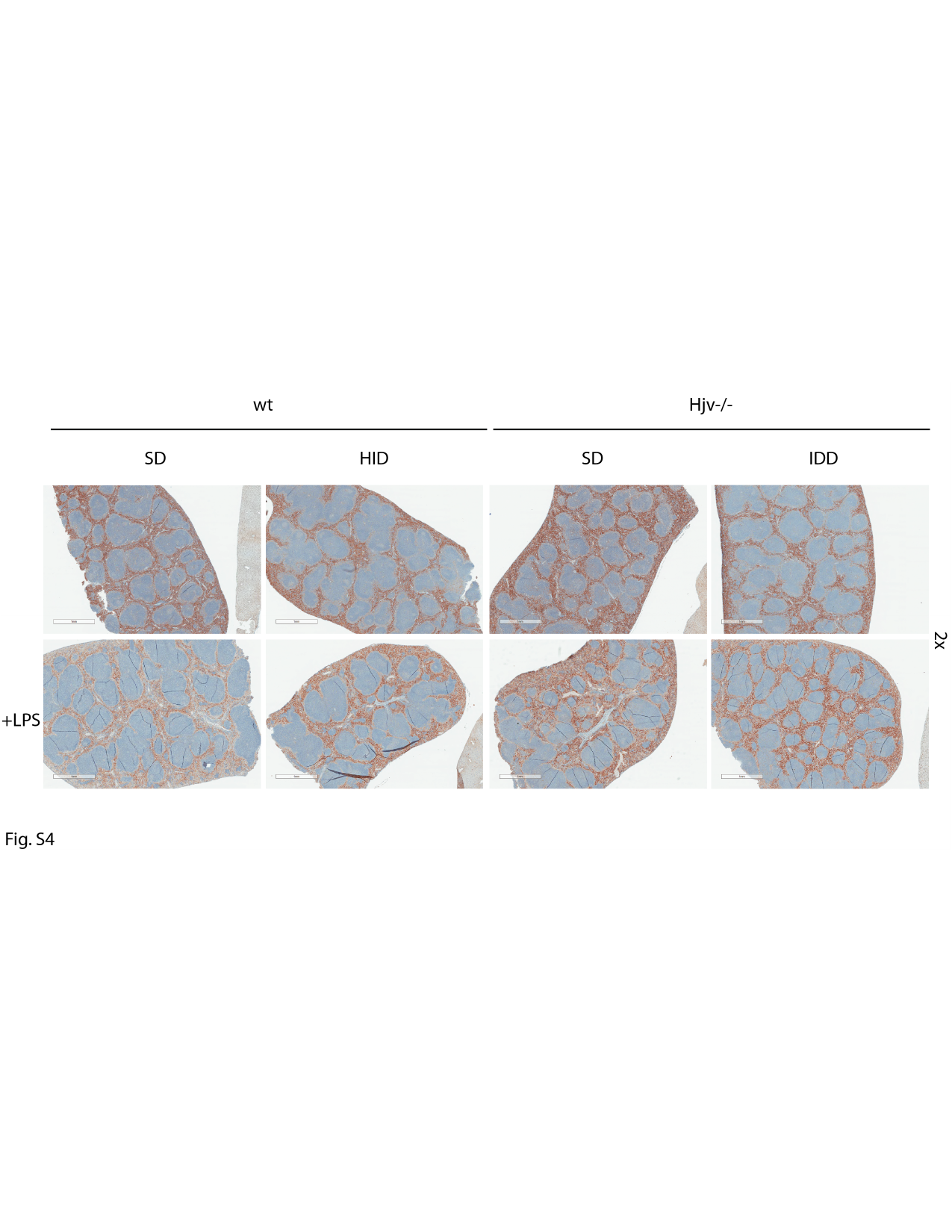

#### Slide 5
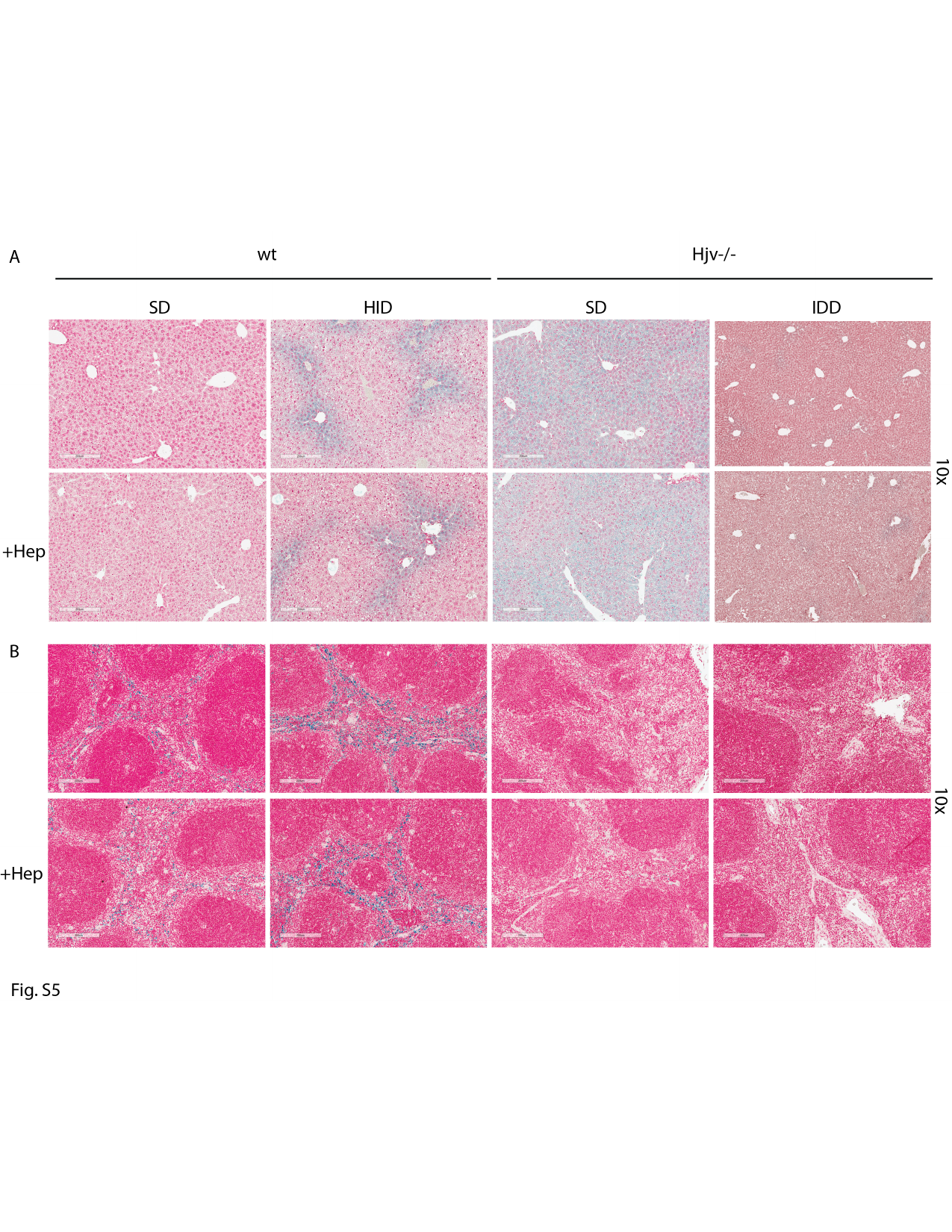

#### Slide 6
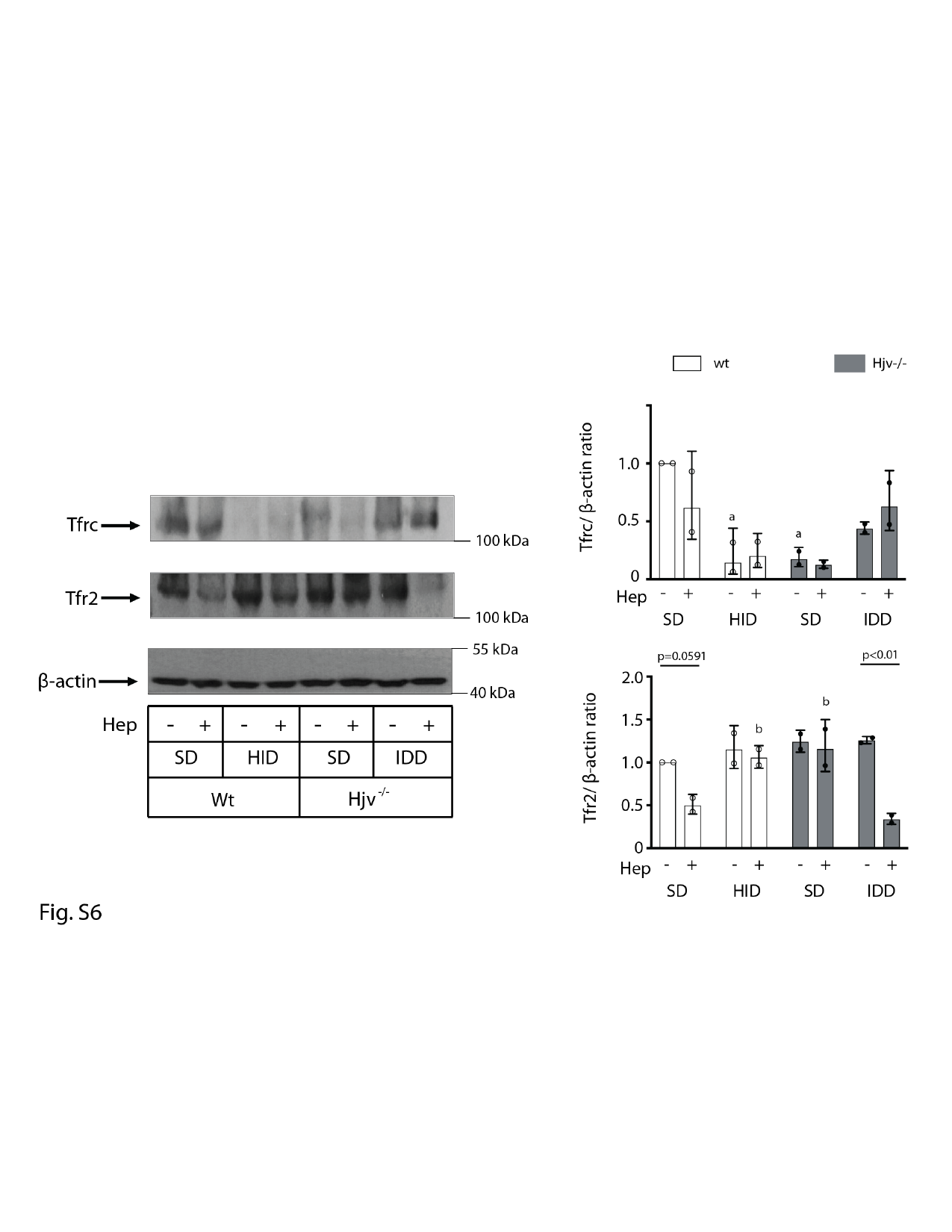

#### Slide 7
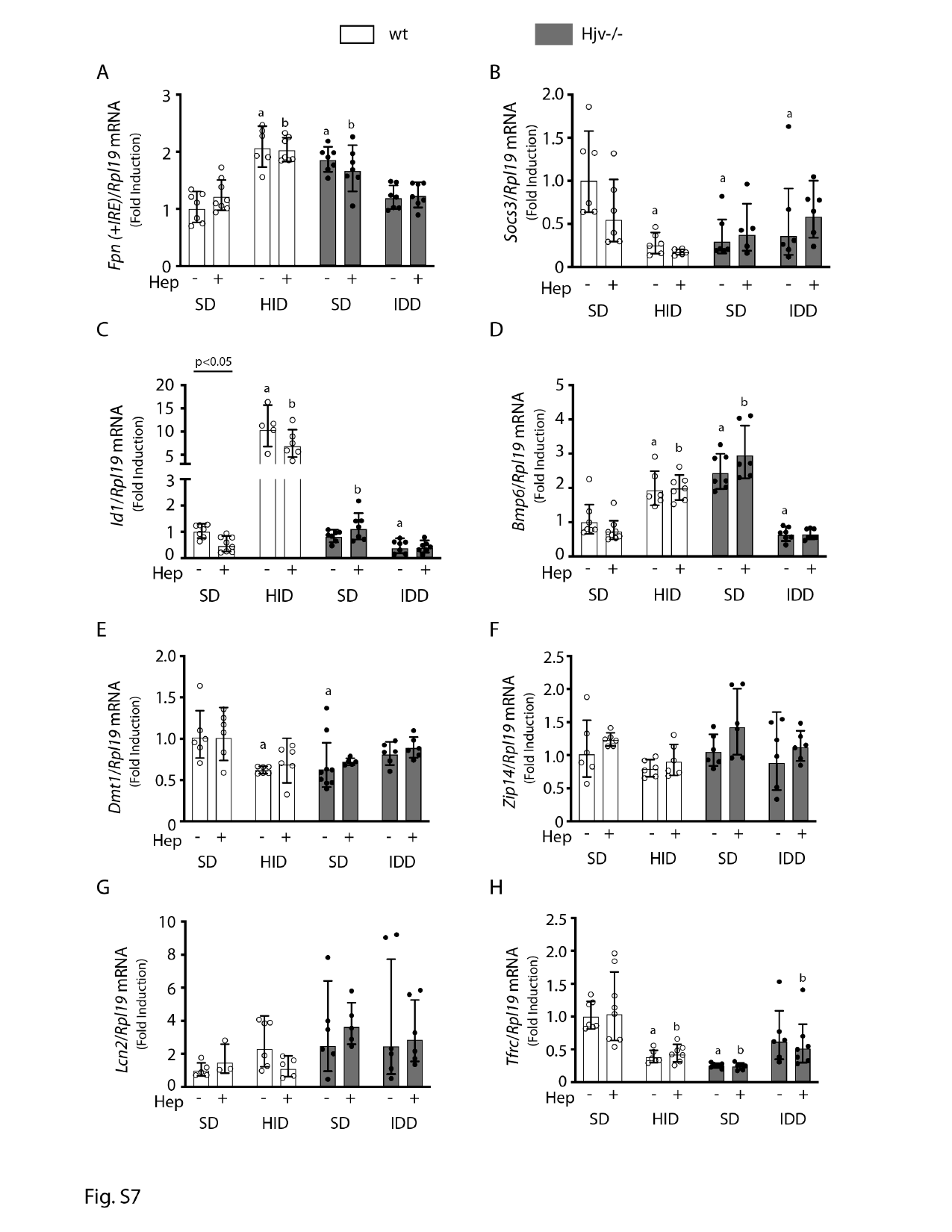

#### Slide 8
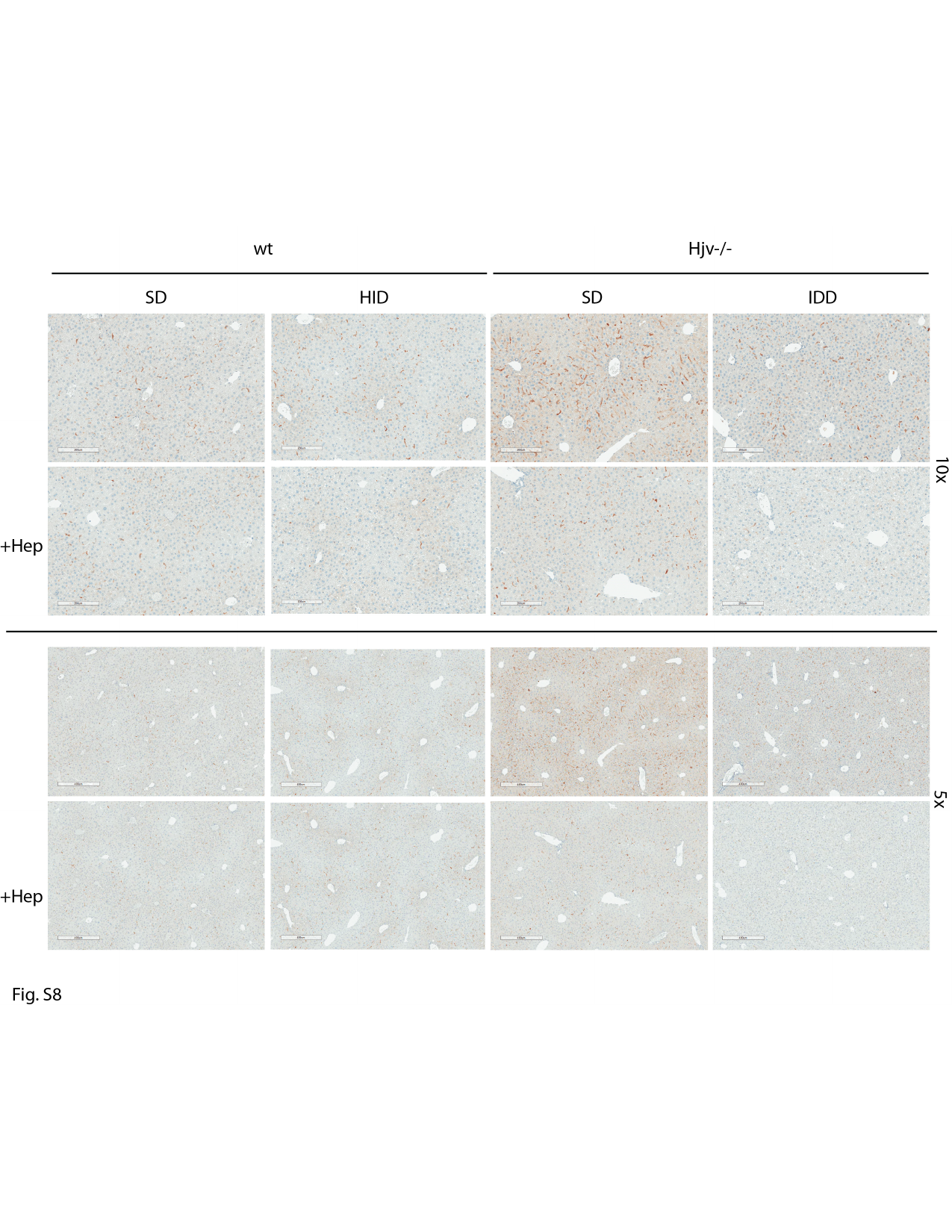

#### Slide 9
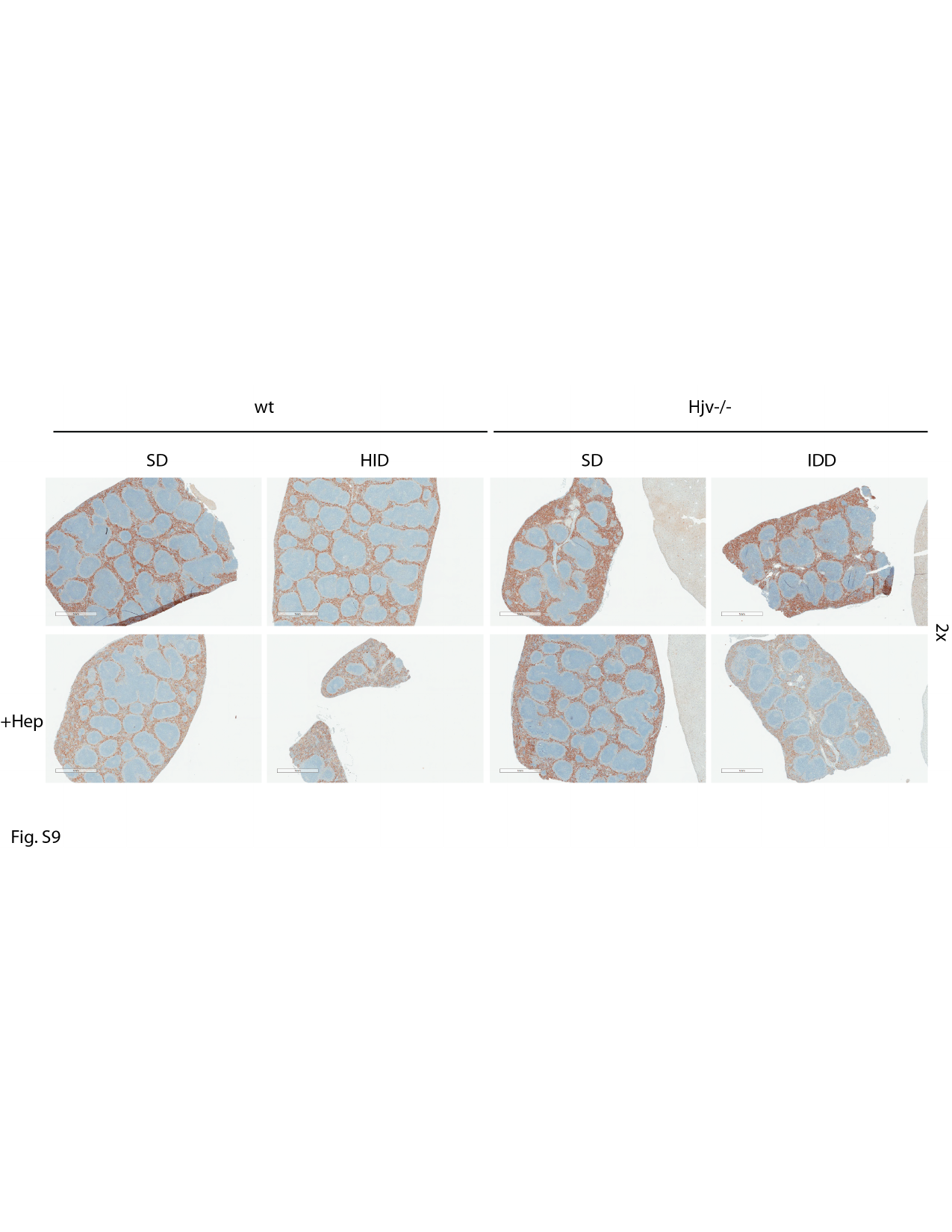
